# *DGCR8* haploinsufficiency leads to primate-specific RNA dysregulation and pluripotency defects

**DOI:** 10.1101/2024.05.02.592145

**Authors:** A Colomer-Boronat, LI Knol, G Peris, L Sanchez, S Peluso, P Tristan-Ramos, A Gazquez-Gutierrez, P Chin, K Gordon, G Barturen, RE Hill, JL Garcia-Perez, A Ivens, S Macias, SR Heras

## Abstract

The 22q11.2 deletion syndrome (22qDS) is caused by a microdeletion in chromosome 22, including *DGCR8*, an essential gene for miRNA production. The contribution of human *DGCR8* hemizygosity to the disease is still unclear. In this study, we generated two human pluripotent cell models containing a single functional *DGCR8* allele to elucidate its role on 22qDS. *DGCR8^+/-^*cells show increased apoptosis as well as self-renewal and differentiation defects in both the naïve and primed states. The expression of primate-specific miRNAs was largely affected, due to impaired miRNA processing and chromatin accessibility. *DGCR8^+/-^* cells also displayed a pronounced reduction in human endogenous retrovirus class H (HERVH) expression, a primate-specific retroelement essential for pluripotency maintenance. Importantly, the reintroduction of primate-specific miRNAs as well as the miR-371-3 cluster rescued the cellular and molecular phenotypes of *DGCR8^+/-^*cells. Our results suggest that *DGCR8* is haploinsufficient in humans and that miRNAs and transposable elements may have co-evolved in primates as part of an essential regulatory network to maintain stem cell identity.

## Introduction

The 22q11.2 deletion syndrome (22qDS) is a human genetic disorder caused by a heterozygous microdeletion at chromosome 22. It is the most common human chromosomal deletion, with an incidence of 1/3,000 to 1/6,000 in live births (McDonald-McGinn *et al*, 2015). Although the deletion is variable in size (1.5Mb to 3Mb), the largest and most frequent deletion (85% patients) affects around 40 protein-coding genes (McDonald-McGinn *et al*., 2015). The major clinical manifestations include developmental disabilities, congenital heart disease, palatal abnormalities, immune deficiency and increased risk of autoimmune diseases and psychiatric illnesses, such as autism and schizophrenia (McDonald-McGinn *et al*., 2015).

The precise relationship between the deletion of specific genes and the subsequent clinical symptoms remains to be fully elucidated. Amongst the 40 genes affected by the microdeletion, the *DGCR8* gene has received much attention, due to its essential role in miRNA biogenesis. MiRNAs are small non-coding RNAs that negatively regulate mRNA stability and translation by imperfect base-pairing to complementary sequences (Gebert & Macrae, 2019). Most miRNAs are transcribed as long primary transcripts (pri-miRNAs) that fold into hairpin structures. These are recognised and cleaved in the nucleus by the Microprocessor complex, which is composed of the dsRNA-binding protein DGCR8 and the RNAse III endonuclease Drosha (Denli *et al*, 2004; Gregory *et al*, 2004; Han *et al*, 2004; Landthaler *et al*, 2004). Next, the excised hairpin is exported to the cytoplasm and further processed by the RNAse III endonuclease Dicer. Finally, one of the strands of the mature miRNA duplex is incorporated into the RNA-induced silencing complex (RISC) to guide repression of the target mRNA (Treiber *et al*, 2019). In addition to miRNA production, the Microprocessor can directly control the levels of mRNAs by cleaving stem-loop structures which resemble pri-miRNAs, including the *DGCR8* transcript itself and retrotransposon-derived RNAs amongst others (Han *et al*, 2009; Heras *et al*, 2013; Knuckles *et al*, 2012; Macias *et al*, 2012; Triboulet *et al*, 2009).

Mouse models of *Dgcr8* heterozygosity have led to contrasting results. While losing one copy of *Dgcr8* in mouse embryonic stem cells (mESCs) was not sufficient to observe significant alterations in the expression of miRNAs by microarrays (Wang *et al*, 2007), analyses of miRNA expression by RT-qPCR or deep sequencing from brain tissue of *Dgcr8* heterozygous mice showed dysregulation of a subgroup of miRNAs (Earls *et al*, 2012; Fenelon *et al*, 2011; Marinaro *et al*, 2017; Schofield *et al*, 2011; Stark *et al*, 2008). These mice also displayed behavioural changes and cognitive defects, which have been attributed to changes in the structure of neuronal dendrites and their synaptic properties (Earls *et al*., 2012; Schofield *et al*., 2011; Stark *et al*., 2008). These studies suggest that the alteration of the structure and function of neuronal circuits in *Dgcr8* heterozygous mice could provide a genetic explanation to the neuropsychiatric manifestations in 22qDS. Although the complete deficiency of *Dgcr8* is embryonically lethal in mice (Wang *et al*., 2007), cell-specific *Dgcr8* ablation has revealed that this gene is also necessary for optimal function of mouse immune cells, including helper T cells, B cells, NK cells and thymic architecture as well as reproductive function, female fertility and spermatogenesis (Bezman *et al*, 2010; Brandl *et al*, 2016; David *et al*, 2011; Khan *et al*, 2014; Kim *et al*, 2016; Zimmermann *et al*, 2014).

Here, we have developed two different human cell models of *DGCR8* heterozygosity, in the embryonic stem cell line H9 and teratocarcinoma cell line PA-1, to investigate if *DGCR8* is haploinsufficient in humans. Our results show that inactivating one *DGCR8* allele results in haploinsufficiency as manifested by dysregulation of miRNA biogenesis but also changes in chromatin accessibility. Despite the functional conservation of *DGCR8* throughout evolution, we found that a high proportion of the affected mature miRNAs are primate specific. As a result, *DGCR8* heterozygote cells display alterations in the gene expression profile associated with pluripotency maintenance and embryonic development, including the human-specific endogenous retrovirus type-H (HERVH) family, a crucial Transposable Element for stem cell identity (Gerdes *et al*, 2016; Lu *et al*, 2014; Römer *et al*, 2017; Wang *et al*, 2014). Interestingly, the alterations caused by *DGCR8^+/-^* are conserved in both the naïve and primed stages of pluripotency. Consistent with these findings, *DGCR8* heterozygote cells display defects in self-renewal and impaired differentiation into the three primary major germ layers. Altogether, these data indicate that *DGCR8* has a significant role in the aetiology of 22qDS that is more relevant than previously was revealed using mouse models of this human genetic disorder.

## Results

### Characterization of *DGCR8* heterozygosity in human pluripotent cellular models

22qDS is caused by a microdeletion in one chromosome 22, resulting in the hemizygosity of around 40 protein coding genes (McDonald-McGinn *et al*., 2015). It is still unclear if the disease originates from the haploinsufficiency of a small subset of these genes or from the absence of the entire region. To investigate this, we assessed if the genes affected by the microdeletion were predicted to be haploinsufficient by comparing their natural variation in the human population using the Genome Aggregation Database (gnomAD) (Karczewski *et al*, 2020). Essential genes are predicted to have a very low frequency of loss-of-function mutations in the general population, as these may be incompatible with life. When the frequency of observed loss-of-function (LoF) mutations is lower than expected (obs/exp ≤ 0.089) the gene is considered haploinsufficient (Karczewski *et al*., 2020). Obs/exp LoF ratios were plotted for each of the genes affected by the most common microdeletion in 22qDS, and only 5 genes were predicted to be haploinsufficient, including *DGCR8* (**Fig 1A**). Similar conclusions were previously reported by (Karbarz, 2020). To validate this prediction, we generated human embryonic stem cells (H9 hESCs) and human embryonic carcinoma cells (PA-1 hECCs), where a single copy of the *DGCR8* gene was inactivated using the CRISPR/Cas9 nickase system. PA-1 cells are a diploid human embryonic teratocarcinoma cell line, which has retained limited pluripotent capacity (Garcia-Perez *et al*, 2010; Zeuthen *et al*, 1980). After targeting, two different *DGCR8* heterozygote clones (HET(1) and HET(2)) for each cell line were selected for further studies (**Fig EV1A-EV1C**). Inactivation of a single *DGCR8* allele resulted in reduced DGCR8 protein expression in both H9 and PA-1 cells, but also of DROSHA, as it requires DGCR8 interaction for stabilisation (Han *et al*., 2009) (**Fig 1B** and **EV1D-EV1F)**. Despite the reduction in DGCR8 expression, H9 HET hESCs did not display obvious changes in colony morphology (**Fig 1C**) or in the expression of the typical pluripotency markers, NANOG and TRA-1-60 (**Fig 1D**). Consistently, no differences were observed in the expression of the other pluripotency markers, OCT4 and SOX2 (**Fig 1E**). Only KLF4 was slightly less abundant in HET H9-HESCs, at both protein and RNA levels (**Fig 1E** and **EV1F-EV1G**). The proportion of alkaline phosphatase expressing colonies was also similar between WT and HET H9-hESCs cells (**Fig EV2A**).

**Figure 1.**
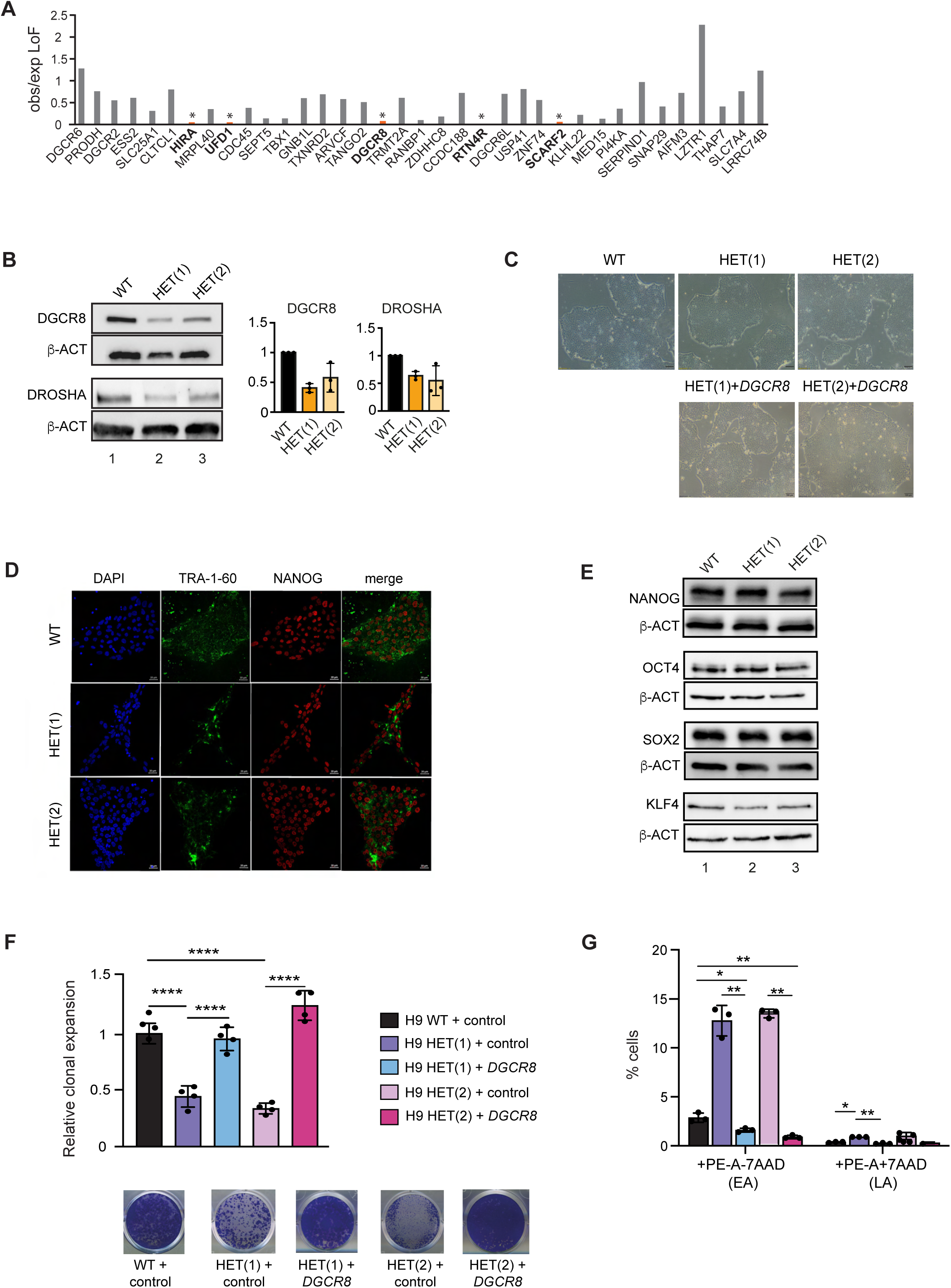
Generation and characterization of *DGCR8* heterozygote cell lines. (**A**) Genes commonly deleted in 22qDS were searched for loss-of-function (LoF) mutations in the human population using the gnomAD. The observed/expected (obs/exp) LoF for each gene was plotted. (*) obs/exp LoF ≤ 0.089, predicted as haploinsufficient. Only genes with at least 10 predicted LoF mutations were considered haploinsufficient (in bold) (**B**) DGCR8 and DROSHA western blot analyses of WT and two different HET *DGCR8* H9 hESC clones. ACTIN serves a loading control. Quantification of DGCR8 and DROSHA protein levels in HET H9 hESCs. Data represents the average of three independent experiments +/− st. dev (**C**) Colony morphology in normal culturing conditions for WT, HET hESCs and HET hESCs with rescued DGCR8 expression by lentiviral transduction (Scale bar = 100 µm) (**D)** Immunofluorescence images for pluripotency markers Tra-1-60 and NANOG (Scale bar = 20 µm) (**E**) NANOG, OCT4, SOX2 and KLF4 western blot analyses of both WT and two different HET *DGCR8* hESC lines. Actin serves as a loading control (**F**) Relative clonal expansion capacity, expressed as the fraction of stained area of HET hESCs lines vs WT transduced with a control-lentiviral vector or with lentivirus expressing human DGCR8. Data represent the average +/− st. dev. of 4 biological replicates. (****) p-val ≤ 0.0001, by one-way ANOVA followed by Tukey’s multiple comparison test (**G**) Percentage of cells in early (EA) and late apoptosis (LA). Data are the average of (n=3) +/− st. dev. (*) p-val ≤ 0.05, (**) p-val ≤ 0.01, by two-way ANOVA followed by Tukey’s multiple comparison test.

In contrast, *DGCR8* HET hESCs displayed a large reduction in their clonal expansion ability when plated at low cell density or as single cells (**Fig 1F** and **EV2B**-**EV2C)** suggesting poor maintenance of self-renewal capacity. Consistent with this finding, both H9 and PA-1 HET showed a decreased doubling time, confirming some proliferation defects (**Fig EV2D)**. Defective proliferation can result from a defect in cell cycle progression and/or increased apoptosis. *DGCR8* HET hESCs displayed delayed cell cycle progression, with a significant accumulation in G0/G1 (**Fig EV2E**), in addition to a significant increase in early (7-AAD negative and PE Annexin V positive) and late (7-AAD positive and PE Annexin V positive) apoptosis (**Fig 1G** and **EV2F**). Importantly, these cellular phenotypes, including defective proliferation, clonal expansion ability, increased apoptosis, and KLF4 misexpression were reverted when DGCR8 expression was rescued in HET cells by lentiviral transduction (**Fig 1F**-**1G, EV1F-EV1G** and **EV2C-EV2D, EV2F**). All these together suggest that inactivation of one *DGCR8* allele in human pluripotent cells results in defective self-renewal capacity, which is characterized by increased apoptosis and cell cycle and proliferation defects.

### *DGCR8* heterozygous hESCs display differentiation defects

To evaluate if *DGCR8* heterozygosity results in defects during human embryonic development, WT and HET *DGCR8* hESCs were differentiated *in vitro* using spontaneous embryoid body (EB) formation. Remarkably, EBs formed by the HET clones were of smaller size in comparison with WT cells, indicating differentiation and proliferation defects (**Fig 2A**). To study the defects in differentiation, pluripotency and differentiation markers’ expression from ectoderm, mesoderm and endoderm, was compared by RT-qPCR at day 0, 7, 14 and 21 of differentiation. WT and HET EB differentiation resulted in a similar repression of the pluripotency markers, *NANOG* and *POU5F1*. The expression of *SOX2* was also similar between WT and HET EBs (**Fig 2B**). Tested ectodermal markers (*OTX2*, *PAX6* and *SOX1*) were also similarly increased during differentiation in both WT and HET hESCs. Only *PAX6* displayed a subtle reduction at day 21 of differentiation (**Fig 2C**). In contrast, the expression of most of the tested mesodermal (*CD34, FOXA2, TBXT* and *HAND1*) and endodermal (*GATA6*, *HNF3*, *SOX7* and *SOX17*) markers was reduced upon differentiation of both clones of HET hESCs, in comparison with WT (**Fig 2D-2E**). These results suggest that *DGCR8* heterozygosity in pluripotent human cells resulted in differentiation defects most markedly for mesodermal and endodermal lineages.

**Figure 2.**
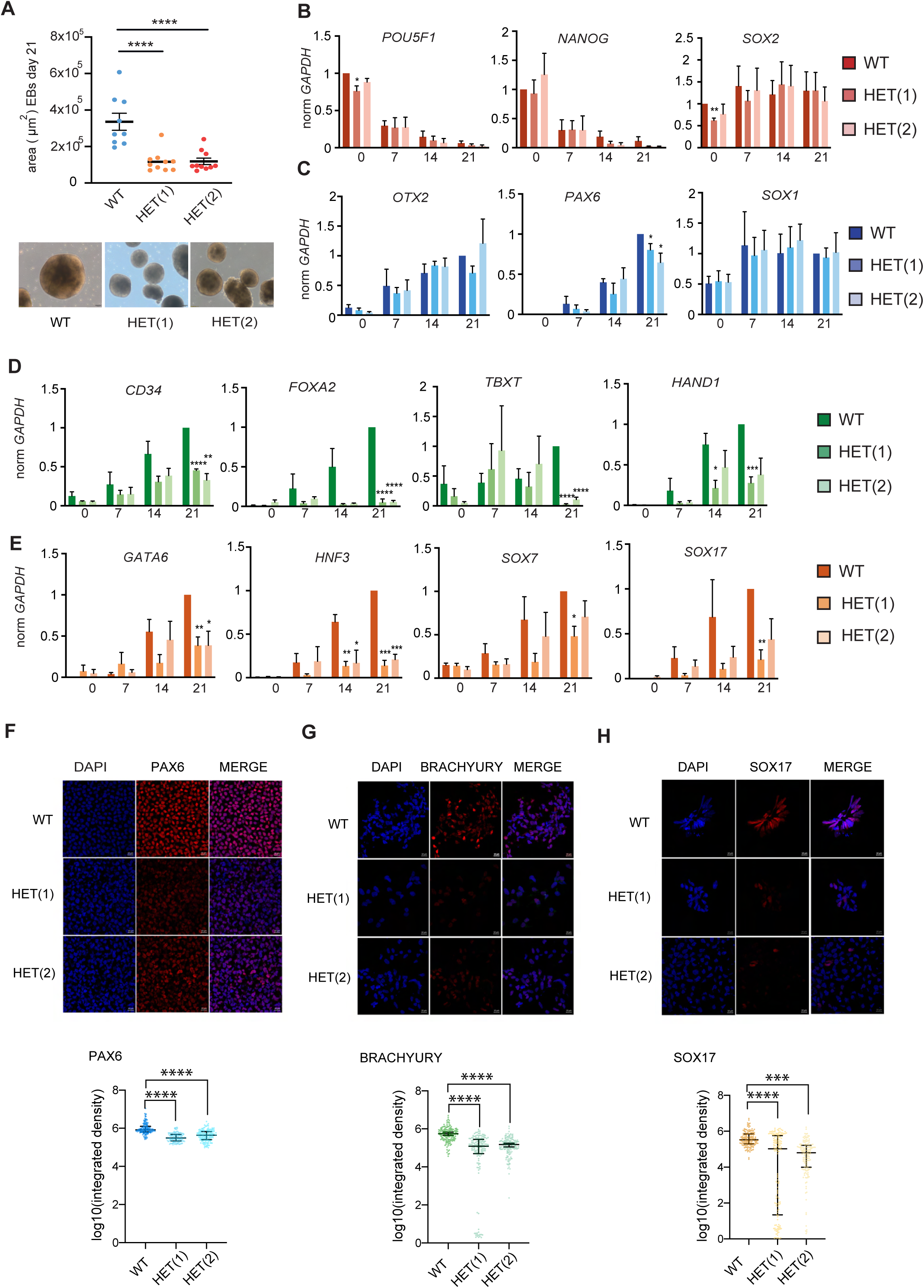
*DGCR8* heterozygote hESCs display differentiation defects. (**A**) Quantification of the area of embryoid bodies at day 21. Data are the average +/− s.e.m; (****) p-val ≤ 0.0001 by generalised linear models with negative binomial distribution, adjusted by Bonferroni post hoc test (top). Representative images of EBs after differentiation at day 21 (bottom) (Scale bar = 100 µm) (**B**) RT-qPCR analyses of pluripotency, (**C**) ectodermal, (**D**) mesodermal and (**E**) endodermal markers after EB differentiation of *DGCR8* HET H9 hESCs compared to the parental cell line (WT). Data are the average (n=3) +/− s.e.m. (*) p-val ≤ 0.5, (**) p-val ≤ 0.01, (***) p-val ≤ 0.001 (****), p-val ≤ 0.0001, by two-tailed Student t-test (**F-H**) Representative immunofluorescence images from WT and HET cells after differentiation to (**F**) ectoderm (PAX6), (**G**) mesoderm (BRACHYURY) and (**H**) endoderm (SOX17). Scale bar = 10 µm. Data represent the log_10_ integrated density for staining in WT and HET cells (n=150), (***) p-val ≤ 0.001, (****) p-val ≤ 0.0001 by generalised linear models with negative binomial distribution, adjusted by Bonferroni post hoc test.

To support these findings, directed differentiation protocols into ectoderm, mesoderm and endoderm were performed followed by immunofluorescence of well-stablished markers for these embryonic layers. This revealed that the average expression of the mesodermal marker BRACHYURY (encoded by the *TBXT*) and the endodermal marker SOX17 was significantly decreased, and nearly absent in a fraction of HET cells. A more homogeneous subtler reduction of the ectodermal marker PAX6 was observed (**Fig 2F-2H**), thus confirming the findings observed upon EB differentiation. All these results suggest that *DGCR8* heterozygosity leads to decreased pluripotency, affecting the establishment of the three major embryonic lineages, with a more pronounced defect in the mesodermal and endodermal germ layers.

### DGCR8 heterozygous hESCs maintain cellular defects in a naïve-like state

Our findings demonstrate that the loss of a functional copy of *DGCR8* results in cellular defects in the biology of hESCs. Conversely, ablation of a single *Dgcr8* allele in mESCs did not lead to significant or clear phenotypes (Wang *et al*., 2007), raising the possibility that the defects associated with human *DGCR8* heterozygosity could be species-specific. Alternatively, these findings could be attributed to the different pluripotency cellular states of hESCs, considered to be in a primed state compared to mESCs, which represent a naïve state (Nichols & Smith, 2009). To rule out this possibility, WT and HET hESCs were induced into a naïve-like state. As a result of this transition, colonies acquired the typical domed morphology of naïve hESCs (**Fig 3A**), preserving the reduced protein levels of both DGCR8 and DROSHA (**Fig 3B**). To confirm successful transition, upregulation of the naïve pluripotency markers *KLF17, DNMT3L, DPPA3* and *DPPA5*, and silencing of the primed pluripotency marker, *DUSP6* was confirmed by qRT-PCR (Guo *et al*, 2017; Messmer *et al*, 2019; Theunissen *et al*, 2014) (**Fig 3C**). Despite the major differences in cellular phenotypes and gene expression profiles of naïve versus primed pluripotent states, the main cellular defects, including increased apoptosis and decreased colony formation capacity, were retained during the naïve stage (**Fig 3D-3G**). All these together suggest that the defects deriving from *DGCR8* heterozygosity may be species-specific rather than cell-state-specific. These results prompted us to further characterise the impact of *DGCR8* heterozygosity at the molecular level.

**Figure 3.**
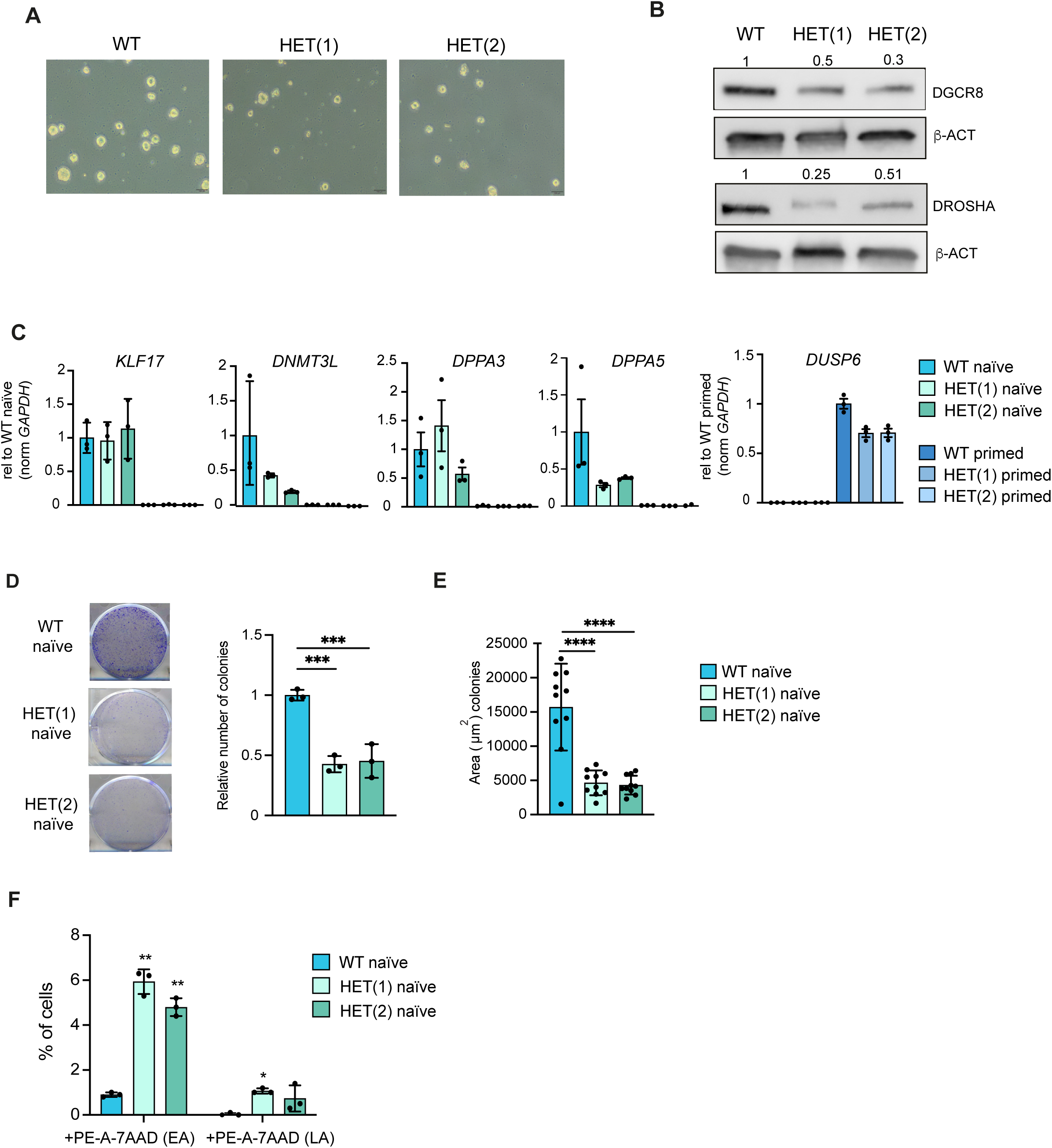
Defects in *DGCR8* heterozygous cells are conserved in a naïve pluripotent state. **(A)** Colony morphology for WT and HET hESCs in naïve culturing conditions (Scale bar = 100 µm) (**B**) DGCR8 and DROSHA western blot analyses of WT and HET *DGCR8* hESC lines. Actin serves as a loading control (**C**) RT-qPCR analyses of naïve (*KLF17*, *DNMT3L*, *DPP3A*, *DPP5A*) and primed (*DUSP6*) markers in both *DGCR8* HET and WT hESCs. Data are normalized to *GAPDH* relative to WT naïve or WT primed, respectively, and represent the average of 3 biological replicates +/− st. dev. (**D**) Relative clonal expansion capacity, expressed as the number of colonies of HET hESCs lines in comparison to WT. Data represents the average (n=3) +/− st. dev. (***) p-val ≤ 0.001, by one-way ANOVA followed by Dunnett’s multiple comparison test (**E**) Quantification of the area in naïve colonies at day 5. Data are the average +/− (n=10) (****) p-val ≤ 0.0001, by one-way ANOVA followed by Dunnett’s multiple comparison test. Colonies are visualised by crystal violet staining (**F**) Percentage of cells in early (EA) and late apoptosis (LA). Data represents the average +/− st. dev of 3 biological replicates. (*) p-val ≤ 0.05, (**) p-val ≤ 0.01 by two-way ANOVA followed by Tukey’s multiple comparison test.

### Expression of primate-specific miRNAs is altered in *DGCR8* heterozygous cells

Considering the defects in proliferation and differentiation of HET cells and the reduced levels of DGCR8 and DROSHA, we next investigated if these phenotypes were linked to abnormal expression of miRNAs. For this purpose, we performed miRNA expression analyses using small RNA-seq of two clones of *DGCR8* HET cells in H9 hESCs and one clone of PA-1 HET cells vs their WT counterparts. *DGCR8* HET hESCs displayed a remarkable reduction in mature miRNAs in a primed stage, with only a small proportion of miRNAs showing a modest upregulation (**Fig 4A**, for complete list of significant differentially expressed miRNAs see **Table EV1**). We also noted that a significant proportion of the common differentially expressed miRNAs in both hESC HET clones were primate-specific (∼30%) (**Fig 4B** and **EV3A**). Dysregulated primate-specific miRNAs mostly belonged to the big miRNA cluster C19MC (Fromm *et al*, 2022) (**Fig 4C**). The expression of the miR-371-3 cluster, homologous to the miR-291-295 cluster in mouse, was also markedly reduced (**Fig 4C**). Interestingly, most members of both clusters share the seed sequence ‘AAGUGC’, which have been previously involved in self-renewal, proliferation, and apoptosis of hESCs (Kobayashi *et al*, 2022; Mong *et al*, 2020; Teijeiro *et al*, 2018). The decreased expression of miRNAs belonging to these clusters was validated by RT-qPCR in both naïve and primed H9 hESC clones and was rescued upon reintroduction of DGCR8 (**Fig 4D-4E**).

**Figure 4.**
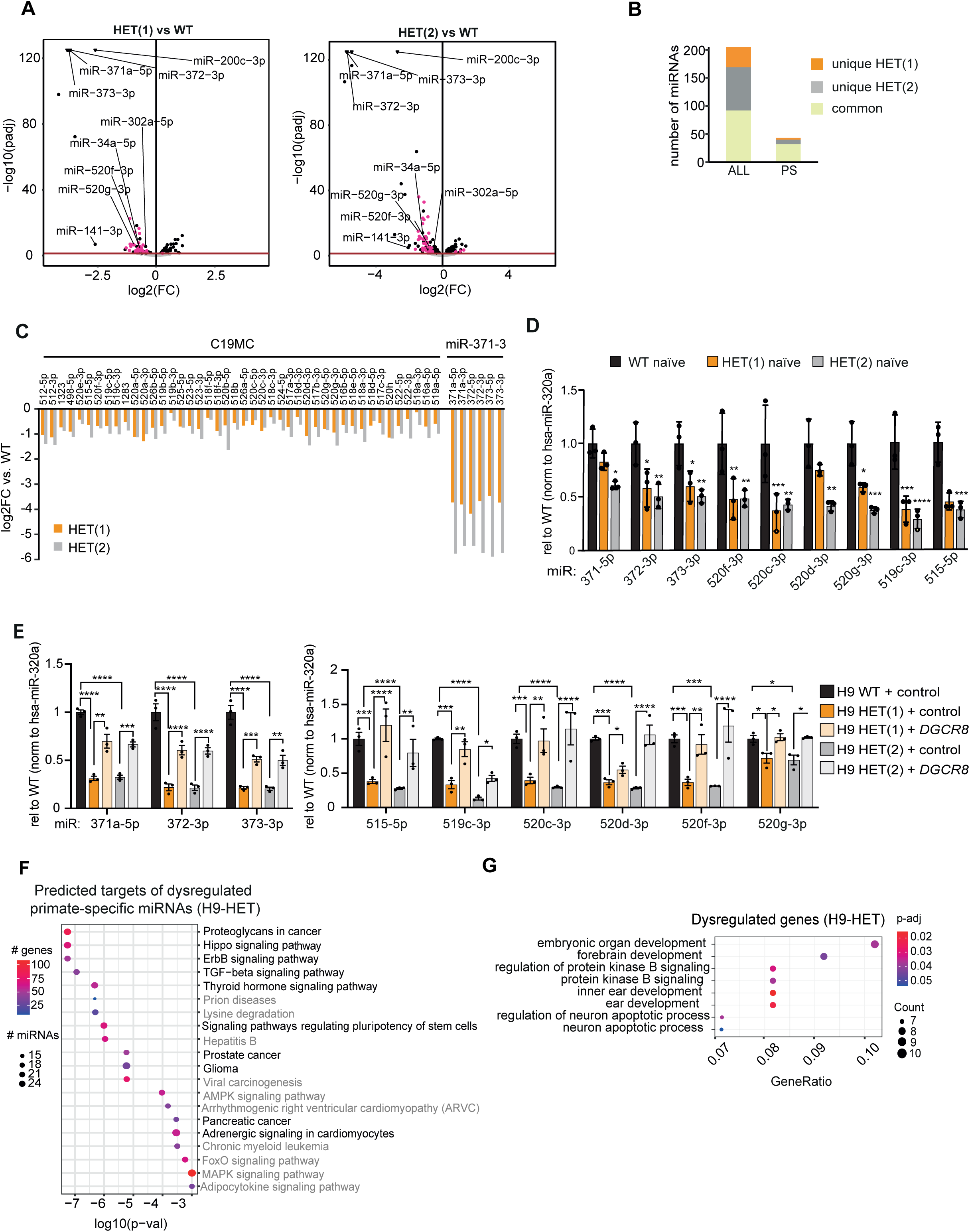
miRNA expression is affected in *DGCR8* heterozygote hESCs. (**A**) Differential expression (log2FC) of miRNAs from two different clones (HET 1 and 2) of *DGCR8* HET hESCs compared to the parental cell line (WT). 69 and 93 miRNAs are downregulated whereas 59 and 76 miRNAs are upregulated in HET(1) and HET(2), respectively. MiRNAs in pink, indicate primate-specific miRNAs (**B**) Commonly dysregulated miRNAs in both HET H9 hESCs are significantly enriched in primate-specific miRNAs (PS) (hypergeometric p-val 1.112e-5) (**C**) Log2FC differential expression of miRNAs derived from the primate-specific C19MC cluster and the miR-317-3 cluster in the two HET clones vs WT H9 hESCs (**D**) RT-qPCR of some mature C19MC and 371-3 miRNAs expression in H9 naïve hESCs and (**E**) in H9 hESCs and HET hESCs transduced with a control or a lentiviral vector expressing human *DGCR8*. Data represent the average (n=3) +/− s.e.m. Expression levels for each miRNAs are normalised to the levels of the DGCR8-independent miRNA, *hsa-miR-320-3p,* and expressed relative to WT levels, (*) p-val ≤ 0.05, (**) p-val ≤ 0.01, (***) p-val ≤ 0.001, (****) p-val ≤ 0.0001, by one-way ANOVA followed by Dunnett’s or Tukey’s multiple comparison test, respectively (**F**) KEGG pathway analyses for the predicted targets (microT-CDS) of the commonly dysregulated primate-specific miRNAs in both HET H9 clones (**G**) GO pathway enrichment for differentially expressed genes in *DGCR8* HET hESCs clones.

To investigate the relevance of dysregulated primate-specific miRNA expression, we performed pathway enrichment analyses with predicted mRNA targets of the primate-specific miRNAs. Obtained pathways included ‘signalling pathways regulating pluripotency of stem cells’ and pathways involved in pluripotency maintenance and self-renewal, as well as embryonic development (e.g., ‘TGF-β signalling’, ‘Hippo signalling’ and ‘ErbB signalling’) (Chan *et al*, 2002; James *et al*, 2005; Ramos & Camargo, 2012; Xu *et al*, 2016) (**Fig 4F**). Enriched pathways for primate-specific miRNAs were similar to those obtained with all the dysregulated miRNAs (10 out of the top 20 predicted pathways), (compare **Fig 4F** and **EV3B,** see common pathways in black). Despite the ontogenic differences between H9-hESCs and PA-1 cells and differences in their miRNA profile, we found that PA-1 HET cells also displayed a similar proportion of miRNAs being differentially expressed, and a similar proportion of those being primate-specific (**Fig EV3A** and **EV3C**). Dysregulated miRNAs were also predicted to regulate pathways involved in pluripotency maintenance and self-renewal (**Fig EV3D** and **EV3E**). These results highlight the potential importance of primate-specific miRNAs, as a subgroup of dysregulated miRNAs during *DGCR8* haploinsufficiency.

### Alterations in the transcriptome of *DGCR8* HET cells

To investigate the impact of miRNA dysregulation on the gene expression programme of *DGCR8* HET cells, we performed total RNA high-throughput sequencing of HET H9 and PA-1 cells (**Fig EV4A** and **EV4B** and **Table EV2**). Functional enrichment analyses of differentially expressed genes (p-adj ≤ 0.05) revealed that affected pathways common to both cell lines were related to development, including ‘embryonic organ development’ (**Fig 4G** and **EV4C**). To understand if changes in gene expression were caused by defective miRNA levels, we investigated the expression of the miRNA targets using the RNA-seq datasets. To this end, we compared the expression of predicted targets for all dysregulated miRNAs (ALL), targets for the subset of primate-specific dysregulated miRNAs (PS), versus non-predicted targets (remaining genes). Changes in the expression of miRNA-predicted targets were significantly different from non-target controls, both for targets of all dysregulated miRNAs and primate-specific miRNAs in PA-1 and H9 HET cells (p-val < 2.22e-16, **Fig EV4D** and **EV4E**). This approach could not differentiate between genes predicted to be regulated by the up-or the down-regulated miRNAs, as many contain predicted binding sites for both of the subgroups. All these findings suggest that, in part, alterations in the gene expression profile of HET cells can be attributed to altered post-transcriptional gene silencing. To better define the molecular mechanisms contributing to the defects of HET cells, we next investigated if the haploinsufficiency resulted in alterations in both well-defined canonical and non-canonical functions of *DGCR8*.

### Heterozygous cells display defects in both the canonical and non-canonical functions of *DGCR8*

To investigate the impact of *DGCR8* heterozygosity on its canonical function, the biogenesis of miRNAs, we quantified Microprocessor cleavage efficiency both *in vitro* and in cells. For *in vitro* purposes, total cell extracts from the three PA-1 cell lines, WT, HET and KO for *DGCR8*, were prepared. Extracts were incubated with radiolabelled pri-miRNAs to visualise precursor miRNA cleavage products as an indirect measurement of processing efficiency. We observed that extracts derived from HET cells only retained partial processing activity when compared to WTs, while KO extracts were not capable of processing pri-miRNAs (**Fig 5A**). Next, to quantify pri-miRNA processing in cells, we measured the Microprocessor Processing Index (MPI) using high-throughput sequencing data of chromatin-associated RNA for WT, HET and KO PA-1 cells, as described in (Conrad *et al*, 2014; Witteveldt *et al*, 2018). For this purpose, cells were fractionated in cytoplasmic, nucleoplasmic and chromatin fractions and confirmed that chromatin was enriched for pri-miRNA transcripts (**EV5A** and **EV5B**). After sequencing, the MPI for each pri-miRNA was calculated as the negative log2 fraction of reads mapping to the hairpin versus reads mapping to the flanks of the primary miRNA. The higher the value, the better processed the pri-miRNA, while values around 0 indicate absence of processing. Both Microprocessor-dependent and independent pri-miRNAs behaved as expected, with accumulation of reads over the hairpin of the Microprocessor-dependent pri-miRNA *pri-miR-374b*, in HET and KO cells, while no changes were observed for the DGCR8-independent miRNA *pri-miR-1234* (**Fig 5B**, for more examples see **EV5C**). We next assessed how the MPI is affected by *DGCR8* heterozygosity and observed that, globally, PA-1 HET cells displayed an intermediate processing efficiency when compared to WT and KO cells (**Fig 5C**, for a full list see **Table EV3**), suggesting that the biogenesis of miRNAs is affected in these cells.

**Figure 5.**
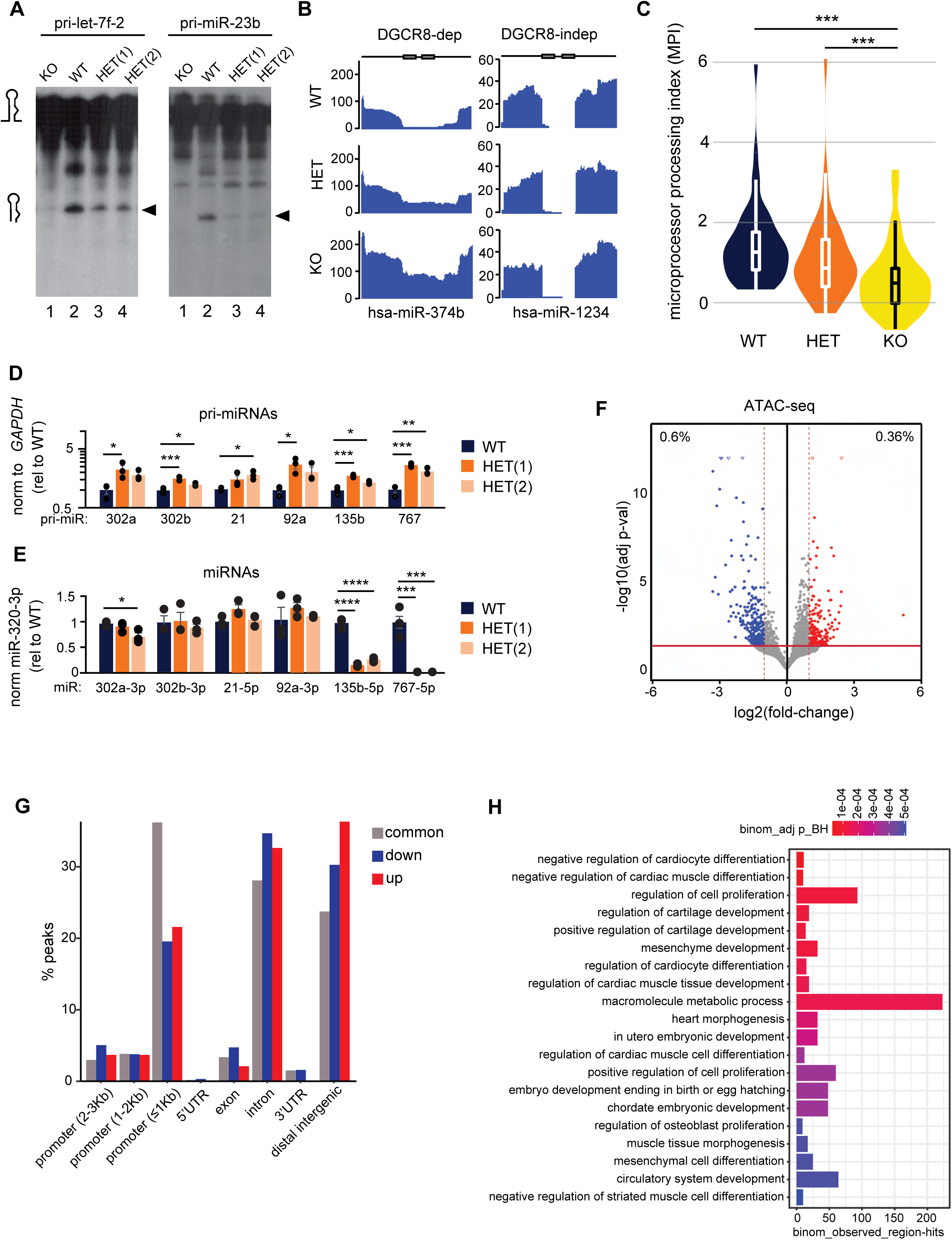
*DGCR8* heterozygosity in PA-1 cells results in impaired miRNA processing and subtle changes on chromatin accessibility. (**A**) *In vitro* processing assays of radiolabelled pri-miRNAs, pri-let-7f-2 and pri-miR-23b, with WT, HET and KO PA-1 derived extracts. Arrow is the resulting cleavage product, the pre-miRNA (**B**) Read depth coverage across a DGCR8-dependent miRNA (*hsa-miR-374b*) and a DGCR8-independent mirtron (*has-miR-1234*) from chromatin-associated RNA high-throughput sequencing in WT, HET and KO PA-1 cells. Grey boxes indicate mature miRNAs, black line represents surrounding genomic regions (**C**) Violin plots for MPI (Microprocessor processing index) values in WT, HET and KO. Only pri-miRNAs with a minimum MPI > 0.3 in WT samples (processed in WT cells) were included in these analyses (***) p-val ≤ 0.001 by one-way ANOVA followed by Tukey’s multiple comparison test (**D**) Quantification of unprocessed pri-miRNA levels by RT-qPCR in WT and HET PA-1 cells. Data are normalised to *GAPDH* and represented relative to WT. Data are the average (n=3) +/− s.e.m. (*) p-val ≤ 0.5, (**) p-val ≤ 0.01, (***) p-val ≤ 0.001 by one-way ANOVA followed by Dunnett’s multiple comparison test (y-axis, logarithmic) (**E**) Quantification of mature miRNA levels by RT-qPCR in WT and HET PA-1 cells. Data are normalised to the DGCR8-independent miRNA, *hsa-miR-320-3p* and relative to WT levels. Data are the average (n=3) +/− s.e.m. (*) p-val ≤ 0.5, (***) p-val ≤ 0.001, (****) p-val ≤ 0.001 by one-way ANOVA followed by Dunnett’s multiple comparison test. (**F**) Differential ATAC accessibility analysis using DEseq2 of HET vs WT PA-1 cells (selected up/down hits with abs. log2FC ≥1, p-adj ≤ 0.05, hits outside plot as open triangles) (**G**) Proportion and genomic distribution of common (grey), less accessible (down, blue), or more accessible ATAC peaks (red, up) in HET vs WT PA-1 (UTR; untranslated region) (**H**) Top twenty more significant GO terms associated with genomic regions that are significantly less accessible in HET cells obtained by rGREAT package.

To explore the relationship between changes in pri-miRNA processing efficiency and mature miRNAs, the levels of several pri-miRNA transcripts and mature miRNAs were compared by RT-qPCR in WT and HET cells. Despite pri-miRNAs accumulation in HET cells, no significant decrease in the mature miRNA levels was observed for some miRNAs, except for miR-135b and miR-767 (**Fig 5D** and **5E**). All these data together suggest that *DGCR8* heterozygosity results in defects in miRNA biogenesis, both *in vitro* and in cells. However, additional mechanisms may contribute to control the final mature miRNA levels.

Besides their canonical role in miRNA biogenesis, DGCR8 and Drosha have also been suggested to regulate gene expression at the transcriptional level, independently of miRNAs. DGCR8 and Drosha have been shown to interact with promoter-proximal regions of human genes enhancing their transcription (Gromak *et al*, 2013). Furthermore, in an indirect manner, DGCR8 has been shown to alter gene transcription by regulating heterochromatin formation through physical association with KAP1 and HP1gamma (Deng *et al*, 2019). Both functions of DGCR8 seem to be independent of the catalytical activity of Drosha. Thus, we next explored whether changes in chromatin structures or accessibility could be associated with the perturbation of gene expression observed in HET cells. For this purpose, we performed ATAC-seq (Assay for Transposase-Accessible Chromatin coupled to high-throughput sequencing) in WT and HET PA-1 cells. As expected, an enrichment of ATAC-seq reads around the transcription start sites (TSS) was observed for both cell lines (see **EV5D)**. Genome-wide differential peak analysis identified 52,347 high-confidence peaks and revealed that there was only a small proportion of peaks that were gained (0.36%, n=190) and approximately twice as many were lost (0.6%, n=317) in HET PA-1 cells (**Fig 5F**). Most differential peaks were located in introns and distal intergenic regions, followed by peaks annotated in proximal (≤ 1 kb) promoters (**Fig 5G**). In agreement with a role of chromatin accessibility in gene expression, 8% of genes containing a differential ATAC peak (+/−10Kb in distance) were also differentially expressed, according to the RNA-seq analysis. Although small, this enrichment was statistically significant (p-val=1.509e-9), suggesting that changes in the chromatin accessibility of HET cells could also be influencing the gene expression profile. To further characterize the functional impact of changes in the accessibility of regulatory regions, we used the rGREAT package which implements the Genomic Regions Enrichment of Annotations Tool (GREAT) (Gu & Hubschmann, 2023). This analysis revealed that some of the most significant terms associated with regions that lost accessibility in HET cells were linked to ‘development’, ‘differentiation’ and “morphogenesis” (**Fig 5H**). All these findings together suggest that the gene expression profile resulting from the loss of a single copy of *DGCR8* could be a combinatorial effect of both miRNA dysregulation and changes in chromatin accessibility.

### *DGCR8* haploinsufficiency reduces expression of primate-restricted endogenous retrovirus HERVH and derived RNAs

Many transposable elements (TEs) are transcribed during early human embryogenesis in a stage-specific manner and their expression is associated with stemness and pluripotency maintenance (Gerdes *et al*., 2016; Torres-Padilla, 2020). For instance, knocking down the RNA derived from the human endogenous retrovirus H (HERVH) or specific HERVH-derived RNAs (e.g., chimeric transcripts driven by their promoter activity) results in the loss of pluripotency and self-renewal capacity of hESCs (Gerdes *et al*., 2016; Lu *et al*., 2014; Römer *et al*., 2017; Wang *et al*., 2014). To investigate if the stemness defects upon *DGCR8* haploinsufficiency originated from aberrant TE expression, we used a pipeline that allows to analyse TE expression at a locus specific level using the RNA-seq datasets from WT and HET H9 hESCs [e.g., Software for Quantifying Interspersed Repeat Elements (SQuIRE) (Yang *et al*, 2019). This analysis revealed that a high number of genomic locations annotated by RepeatMasker as endogenous retroviruses type 1 (ERV1) were significantly downregulated in both H9 hESC HET clones compared to WT cells (log2FC < –1; p-adj ≤ 0.05; **Fig 6A**). ERV1 retrotransposons have a structure resembling simple retroviruses, as they encode for *gag* and *pol* genes, and are flanked by ∼450bp Long Terminal Repeats (LTRs). However, genomic analysis indicate that these retrotransposons are not currently active in the human genome (Gerdes *et al*., 2016). Downregulated ERV1 loci from HET H9 hESCs mostly belonged to the family members of the primate-specific endogenous retrovirus class H, HERVH, with reads mapping to both their internal region (HERVH-int) and LTRs (known as LTR7) (89.2% and 70.49% of the ERV1 mapped reads in HET1 and HET2, respectively) (**Fig 6B**, for locus-specific examples see **Fig 6C**, upper panels).

**Figure 6.**
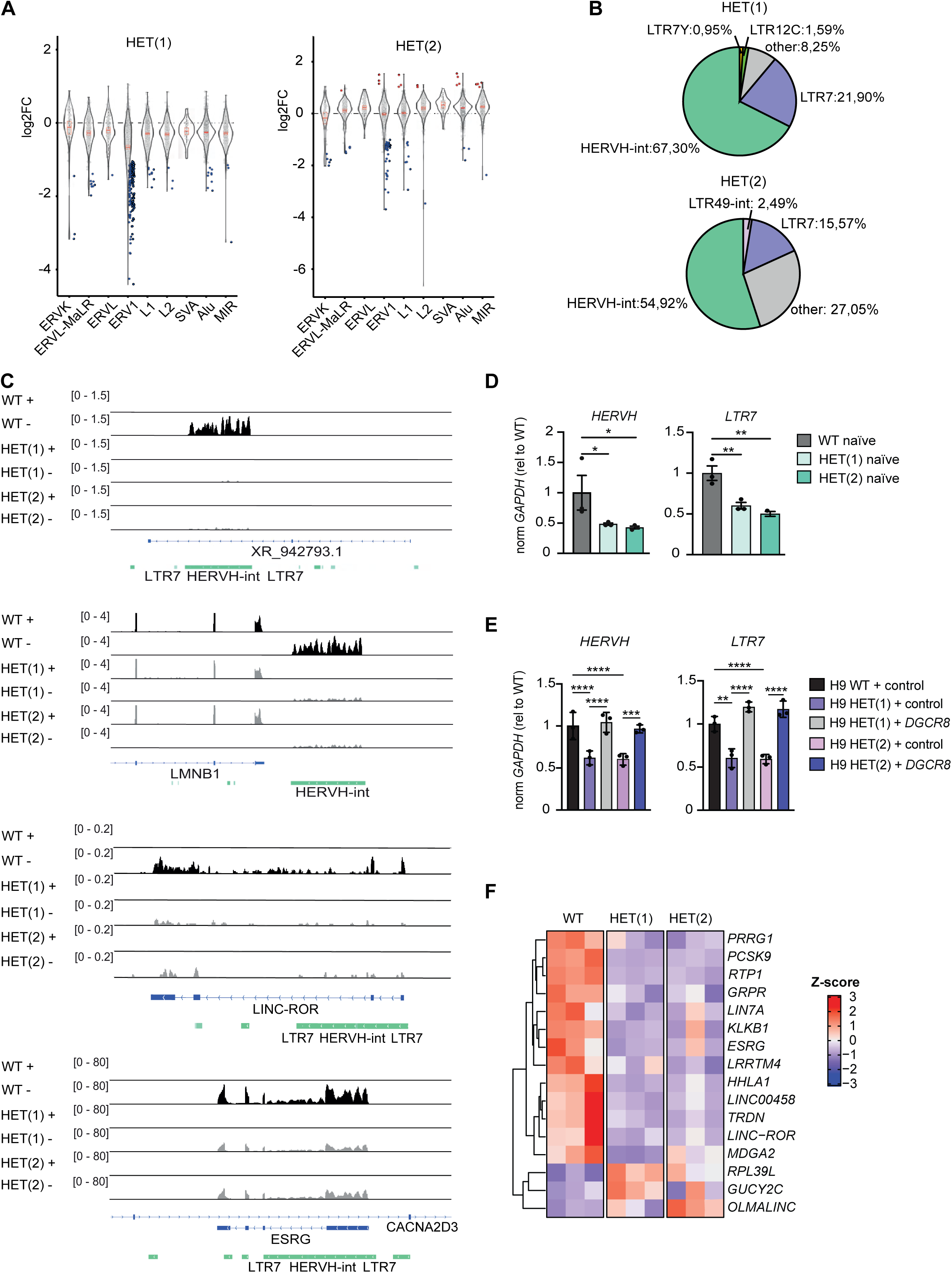
DGCR8*-*mediated control of HERVH expression. **(A)** Violin plots for locus-specific expression of retrotransposons in the two *DGCR8* hESC clones compared to WT cells. Significantly downregulated (log2FC < –1; p-val ≤ 0.05; baseMean > 100) and upregulated (log2FC > 1; p-val ≤ 0.05; baseMean > 100) elements are represented as blue and red dots, respectively (**B**) Distribution of downregulated ERV1 subfamilies in both HET hESC clones (**C**) Genome browser view of RNA-seq data from WT and HET *DGCR8* hESCs clones. Two representative regions containing differentially expressed LTR7-HERVH elements are shown, *XR_942793.1* and *LMNB1* (top) and two chimeric transcripts driven by LTR7 promoter activity are shown as representative examples, *LINC-ROR* and *ESRG* (bottom). Sense (+) and antisense (-) strands are represented. Genes are depicted in blue and LTR7-HERVH in green (**D**) RT-qPCR analyses for HERVH-int and LTR7 in WT and HET naïve hESCs (**E**) the same as (**D**), but in WT and HET primed hESCs transduced with lentiviral control vector or with lentivirus expressing *DGCR8.* Data are normalised to *GAPDH* and relative to WT levels. Data are the average (n=3) +/− st. dev. (*) p-val ≤ 0.05, (**) p-val ≤ 0.01, (***) p-val ≤ 0.001, (****) p-val ≤ 0.0001, by one-way ANOVA, followed by Tukey’s or Dunnett’s multiple comparison test, respectively (**F**) Heatmap for differentially expressed (p-adj ≤ 0.05) chimeric transcripts and lncRNAs driven by LTR7 promoter activity in HET hESCs.

HERVH elements are typically expressed in pluripotent human cells. Indeed, nearly half of all HERVH genomic copies (550 out of the 1225 full-length HERVH copies) are transcribed in hESC, although only a relatively small subset of loci (n∼117) is highly expressed (Wang *et al*., 2014). High expression of HERVHs in hESCs is mostly driven by LTR7 rather than its counterparts, LTR7b, LTR7c or LTR7y (Wang *et al*., 2014). Remarkably, a significant proportion of the downregulated HERVH elements in both HET H9 hESCs clones belonged to the subgroup previously shown to be highly expressed in hESCs and mostly associated with LTR7 promoter activity (formerly known as Type I subfamily) (**Table EV4**). To validate if HERVH transcripts were reduced in H9 hESC HET cells, we measured the expression of both, total HERVH RNA levels and specific HERVH-derived transcripts. Using RT-qPCR, we observed a global reduction in LTR7 and HERVH-int mRNAs in both naïve and primed *DGCR8* hESCs HET, which were rescued upon reintroduction of DGCR8 (**Fig 6D** and **6E**). Additionally, we observed a downregulation of specific HERVH-derived transcripts, as defined by (Wang *et al*., 2014), including the hESCs-specific lncRNAs and chimeric transcripts (**Fig 6C, lower panels,** and **6F**). Downregulated HERVH-derived transcripts, including *linc-ROR, ESRG* and *LINC00458*, have also been previously associated with pluripotency maintenance (Römer *et al*., 2017; Sexton *et al*, 2022).

Thus, these data strongly suggest that *DGCR8* haploinsufficiency in pluripotent human cells lead to misregulation of HERVH expression. Furthermore, these data further support that the phenotype associated with *DGCR8* heterozygosity is species-specific, and that in human cells impacts expression of primate-specific miRNAs and HERVH retroelements.

### C19MC and miR-371-373 miRNAs restore the molecular and cellular phenotype of *DGCR8* heterozygous cells

Our results suggest that *DGCR8* HET pluripotent human cells display two different primate-specific molecular phenotypes. First, the downregulation of primate-specific miRNAs and second, HERVH derived transcripts, both of which are necessary for embryogenesis and pluripotency maintenance. We next wanted to test if HERVH downregulation was a consequence of the depletion of certain miRNAs, or if miRNAs and TEs were operating on separate pathways. To this end, HET cells were transfected with two different pools of miRNAs to test their ability to rescue HERVH expression. Full restoration of HERVH RNA levels was observed after reintroduction of miRNAs belonging to the miR-371-3 cluster (hsa-miR-372-3p and hsa-miR-373-3p), and a partial rescue was observed after transfection of four miRNAs from C19MC cluster (hsa-miR-520g-3p, hsa-miR-520d-3p, hsa-miR-519c-3p, hsa-miR-515-5p) (**Fig 7A**). Notably, the protein levels of KLF4 were also restored upon overexpression of miRNAs from both clusters (**Fig 7B**). Next, we assessed whether overexpression of those miRNAs was also sufficient to restore some of the cellular phenotypes of HET cells. Indeed, transient transfection with miRNAs from both clusters, either as single miRNAs or as pools, re-established the clonal expansion capacity of *DGCR8* HET cells (**Fig 7C** and **7D**).

**Figure 7.**
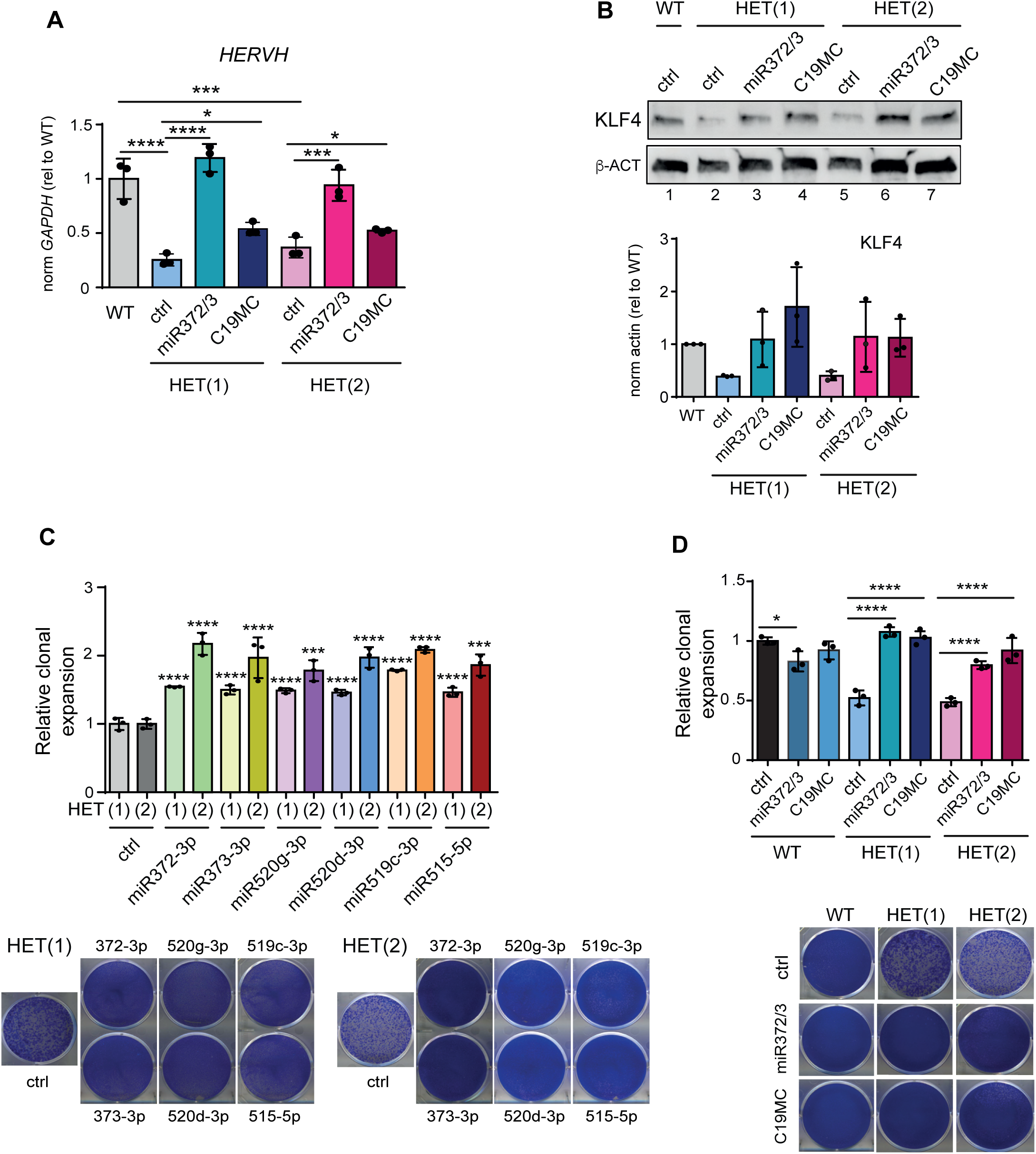
C19MC and miR-371-373 miRNAs rescue molecular and cellular defects in HET hESCs. (**A**) RT-qPCR for HERVH-int in WT and HET hESCs transfected with mimic control or with two miRNA mimics belonging to 371-3 cluster (372-3p and 373-3p) or five miRNA mimics from C19MC cluster (520g-3p, 520d-3p, 519c-3p and 515-5p). Data are normalised to *GAPDH* and relative to WT levels. Data are the average (n=3) +/− st. dev. (*) p-val ≤ 0.05, (***) p-val ≤ 0.001, (****) p-val ≤ 0.0001, by one-way ANOVA, followed by Sidak’s multiple comparison test (**B**) KLF4 western-blot analyses for WT and HET hESCs transfected with mimic control or with miRNA mimics belonging to 371-3 cluster or miRNA mimics from C19MC cluster. Actin serves as a loading control (top). Quantification of KLF4 protein levels in WT and HETs cells transfected with miRNAs mimics. Data are the average (n=3) +/− st. dev and relative to WT levels (down) (**C**) Relative clonal expansion capacity, expressed as fraction of stained area of HET hESCs transfected with miRNA mimics relative to HET hESCs lines transfected with a control miRNA mimic. Data represent the average +/− st. dev. of 3 biological replicates. (***) p-val ≤ 0.001, (****) p-val ≤ 0.0001 by one-way ANOVA followed by Dunnett’s multiple comparison test (**D**) Relative clonal expansion capacity, expressed as fraction of stained area of WT and HET hESCs transfected with a control miRNA mimic or a pool of miRNA mimics from miR-371-3 cluster (372-3p and 373-3p) or with C19MC miRNA mimics (520d-3p, 520g-3p, 519c-3p and 515-5p) and relative to WT transfected with the control mimic. Data represent the average +/− st. dev. of 3 biological replicates. (*) p-val ≤ 0.05, (****) p-val ≤ 0.0001 by one-way ANOVA, followed by Sidak’s, multiple comparison test. Colonies are visualised by crystal violet staining.

These findings indicate that the primate-specific miRNA cluster C19MC, along with the miR-371-373 cluster, play crucial roles in human embryonic stem cell maintenance. Also, we showed that decreased KLF4 and HERVH RNA levels are a consequence of miRNA dysregulation. However, we predict that all these factors are acting in concert and contribute to the observed pluripotency defects of *DGCR8* HET cells.

## Discussion

In this study, we have used two independent human pluripotent cellular models containing a single functional *DGCR8* allele to understand its relevance in the context of 22qDS. Our results indicate that *DGCR8* is haploinsufficient and that the most prominent defects are primate-specific. Previous attempts to study the consequences of *DGCR8* haploinsufficiency were performed in mice and led to conflicting conclusions. Mouse ESCs harbouring a single *Dgcr8* gene do not show significant defects in miRNA levels or differentiation (Han *et al*., 2009; Wang *et al*., 2007). Despite the apparently negligible consequences, *Dgcr8* HET mice displayed behavioural and neuronal defects, which were attributed to altered miRNA expression (Stark *et al*., 2008). In contrast to some of these findings in mice, our human models showed defects in pluripotency and dysregulation of miRNA expression. Inconsistencies between human and mouse models were also previously highlighted when comparing the transcriptome of 22qDS-derived neurons and those derived from the mouse model (*Df16A^+/−^*), where no overlap was found (Khan *et al*, 2020; Sun *et al*, 2018). We hypothesise that this discrepancy is due to intrinsic differences between species. For instance, our results showed that *DGCR8* HET hESCs display increased apoptosis and defects in self-renewal, which demonstrates the importance of *DGCR8* for hESC survival. In agreement with this finding, *DGCR8* has also been identified as an essential gene for the survival of haploid hESCs (Yilmaz *et al*, 2018). In contrast, the total absence of *Dgcr8* in mESCs results in a reduction in proliferation but without changes in cell death or self-renewal ability (Wang *et al*., 2007). Similarly, species-driven differences have been observed for DICER deficiency, another essential factor for miRNA biogenesis. While hESCs require *DICER1* for self-renewal, it seems to be dispensable for mESC survival (Teijeiro *et al*., 2018). These discrepancies could arise from differential sensitivities to unbalanced miRNA levels within the different developmental stages that mouse and human ESCs represent (naïve versus primed states, respectively). However, our results showed that hESCs maintain the main molecular and cellular defects after induction into a naïve-like stage, arguing against a cell-state-specific phenomena and supporting species-specific differences. In agreement with these findings, our results suggest that primate-specific miRNA dysregulation could be largely responsible for the stemness defects in *DGCR8* HET hESCs. We observed that a third of the miRNAs that are affected by *DGCR8* haploinsufficiency are primate-specific and not present in rodents, some of which have been shown to be associated with pluripotency maintenance and self-renewal, including the C19MC cluster (Lin *et al*, 2010; Mong *et al*., 2020). Ree et al has also reported defects in C19MC miRNA processing, despite an inconsistent efficiency in targeting *DGCR8* expression in human cells (Reé *et al*, 2022). The clonal expansion defects observed in *DGCR8* HET hESCs could be rescued by reintroducing four independent miRNAs belonging to this cluster, indicating its contribution to proliferation. Interestingly, only two of the rescued miRNAs, miR-520d-3p and miR-519c-3p, share the seed sequence with miR-371-373 suggesting that the functions of these clusters in human pluripotent cells are partially redundant. Similar to our findings, Teijero et al. showed that reintroduction of miR-372-3p and miR-373-3p rescues the increased apoptosis of *DICER* knockout hESCs (Teijeiro *et al*., 2018). Importantly, human and mouse early development display remarkable differences, some of which seem to be driven by species-specific expression of transposable elements (Gerdes *et al*., 2016). For instance, the murine endogenous retrovirus-L (MERVL) is transiently upregulated at the two-cell stage and is essential for mouse preimplantation development (Sakashita *et al*, 2023). The endogenous retrovirus that colonised the common ancestor of apes, HERVH, is highly expressed in human pluripotent stem cells (hESCs and iPSCs) and epiblast, where it appears to play a role in promoting self-renewal and pluripotency (Goke *et al*, 2015; Grow *et al*, 2015; Wang *et al*., 2014). Notably, almost all HERVH elements expressed in hESCs belong to a subfamily of elements transcribed from LTR7, which contains binding sites for specific transcription factors, including KLF4 (Carter *et al*, 2022; Ohnuki *et al*, 2014). Haploinsufficiency of *DGCR8* led to a significant reduction of the HERVH/LTR7 subfamily transcripts and KLF4 levels, and these were both restored upon reintroduction of miRNAs. Our findings suggest that HERVH/LTR7 silencing is a consequence of miRNA downregulation and could be potentially mediated by KLF4 knockdown. The reduction of this primate-specific endogenous retrovirus, in both naïve and primed stages, may also contribute to some of the cellular phenotypes of *DGCR8* HET hESCs. These findings lead us to hypothesise that important primate-specific non-coding RNAs, including those derived from miRNAs and transposable elements, may have co-evolved to orchestrate crucial aspects of pluripotency maintenance in primates.

Our previous results showed that RNAs derived from other types of TEs (LINEs and SINEs) are bound and processed by the Microprocessor (Heras *et al*., 2013). However, HET hESCs showed no significant differences in the expression levels of these active retrotransposons, suggesting that inactivating a single allele of *DGCR8* is not sufficient to abolish the post-transcriptional control of LINE/SINE in pluripotent cells.

To identify which other functions of DGCR8 were altered in HET cells, we studied both the canonical and less known functions. Similar to Stark et al. findings in mice (Stark *et al*., 2008), human *DGCR8* HET cells showed differential expression of a small proportion of miRNAs. However, miRNA biogenesis defects did not seem to be directly correlated with mature miRNA levels, indicating that additional factors may influence mature miRNA abundance, including differences in the transcription of precursor pri-miRNAs. In addition to miRNA biogenesis, DGCR8 has been implicated in stimulating RNA-pol II transcription, but also promoting heterochromatin formation through interaction with KAP1 (Deng *et al*., 2019; Gromak *et al*., 2013). ATAC-seq analysis of *DGCR8* HET cells only showed subtle changes in chromatin accessibility. Interestingly, affected regions were associated with development pathways, suggesting that changes at chromatin level could also be involved in some of the cellular phenotypes characterised in HET cells.

Collectively, we show that DGCR8 results in haploinsufficiency by altering the expression of primate-specific miRNAs, as well as of primate-specific transposable elements in human pluripotent cells independently of the developmental cellular stage. These findings stress the potential limitations of studying the function of human genes in evolutionary distant animal models (e.g., rodents, zebrafish, etc) where a proportion of the genome, especially the non-coding genome, is only partially conserved. Particularly, our results suggest that *DGCR8* could have a more profound role in the developmental issues present in 22qDS to what it had been previously suggested using mouse models and open new avenues to design novel targets for rescuing some of the developmental defects in 22qDS patients.

## Material and methods

### Cell lines

All cell lines were grown at 37℃ and 5% CO_2_. H9 hESCs were obtained from WiCell and cultured in mTeSR1 media (STEMCELL Technologies) in plates coated with Matrigel (Corning). Human PA-1 cells were cultured in MEM (Gibco) supplemented with GlutaMAX, 20% heat-inactivated FBS (Gibco), 100 U/mL penicillin-streptomycin (P/S, Invitrogen) and 0.1 mM Non-Essential Amino Acids (Gibco). iROCK (10 uM, Y-27632, Sigma) was added to medium during the first 24 hours after splitting, followed by replacement with fresh media. HEK293T cells were obtained from ATCC and cultured in high-glucose DMEM (Gibco) supplemented with GlutaMAX, 10% FBS, (Hyclone) and 100 U/mL P/S. STR (Short tandem repeat) analysis was carried out at the Genomic Unit (Genyo, Granada).

### Generation of CRISPR-edited clonal cell lines

Guide RNAs (gRNA) A (GCACCACTGGACGTTTGCAG) and B (GAGGTAATGGACGTTGGCTC) were designed to target exon 2 of *DGCR8*, after the start codon, using the double nickase design from CRISPR Design Tool (http://tools.genome-engineering.org). gRNAs were cloned into pX461 (pSpCas9n(BB)-2A-GFP, Addgene ID # 48140) as in (Ran *et al*, 2013). For CRISPR targeting, H9 hESCs were maintained in E8 media (DMEM/F12, L-ascorbic acid-2-phosphate magnesium (64 mg/l), sodium selenium (14 µg/l), FGF2 (100 µg/l), insulin (19.4 mg/l), sodium bicarbonate (1.74 g/l), NaCl (5 mM) and holo-transferrin (10.6 mg/l) and TGFβ1 (1.8 µg/l)). Approximately, 2×10^6^ hESCs were nucleofected with 2 µg each of pX461-sgRNA(A) and (B) using the V-Kit solution (Amaxa) and the A-23 program and seeded at a density of 2×10^3^ in a Matrigel-coated plate, as in (Macia *et al*, 2017). Control transfection with plasmid pMAX-EGFP (Amaxa) revealed that ∼50% of cells were GFP+ by fluorescence microscopy. Five days after nucleofection, cells were dissociated with TrypLE and single cell clonal cell lines generated by limited dilution in 96-well coated plates.

iROCK inhibitor was added to the media during passaging to increase cell survival. After two passages, a fraction of the cells was used for genomic DNA extraction using QuickExtract DNA Solution (Lucigen). PCR from genomic DNA was performed using KAPA2G Fast Hotstart Ready Mix PCR Kit (Kapa Biosystems). PCR products were cloned in pGEM-T Easy Vector (Promega), and at least 10 clones were sent for Sanger sequencing for each cell line. Two *DGCR8*^+/−^ (HET) clones with a frameshift mutation in one allele were selected for further studies. These clones were named H9 hESCs HET(1) and HET(2). For CRISPR targeting of PA-1 cells, 1.25 µg of each pX461-sgRNA(A) and (B) were co-transfected using Lipofectamine 2000. GFP^+^ cells were sorted 48h post-transfection using a FACSAria Cell Sorter (BD) and seeded in 96-well plates. iROCK was added during passaging. Genomic DNA sequencing of PA-1 clones was performed as described for H9 hESC clones. As no heterozygote clones were obtained during the first round of targeting, one clone containing frameshift mutations in both *DGCR8* alleles (*DGCR8^-/-^* KO) was used to generate *DGCR8^+/−^* (HET) cells by repairing one of the mutated alleles. To this end, 245 pmols of crRNA (AGGTAATGGACGTTGGACGT), complementary to only one of the mutated alleles in the KO cells, and 245 pmols of tracrRNA (IDT) were incubated in 25 µl nuclease free buffer for 5 min at 95℃ and allowed to anneal at room temperature (RT). The resulting gRNA was incubated with 25 µg of Cas9 protein (IDT) for 15-25 min at 37℃ prior to transfection. 1.2 x 10^6^ PA-1 *DGCR8^-/-^* (KO) cells were resuspended in 80 µl of buffer T (Neon Transfection system), and the Cas9/gRNA mixture and 300 pmol of the repair template were added to be electroporated using 3 pulses of 1600V and 10 milliseconds. Single cell clones were obtained by limiting dilution in 96-well plates. To test successful gene editing, genomic DNA was extracted by incubating cells in lysis buffer (30 mM Tris-HCl pH 8.0, 10 mM EDTA, 0.1% SDS, 0.5% Tween-20 and 10 ug/ml Proteinase K) for 15 min at RT. Next, lysate was transferred to 57℃ for 10 min followed by 98℃ for 10 min. From this mix, amplification of the sequence of interest was performed by PCR, and products were cloned in pGEM-T Easy Vector. Successful gene editing was confirmed by Sanger sequencing. Two clones were selected for further studies and named PA-1 HET(1) and HET(2). All primers used are listed in **Table 1**.

**Table 1.**
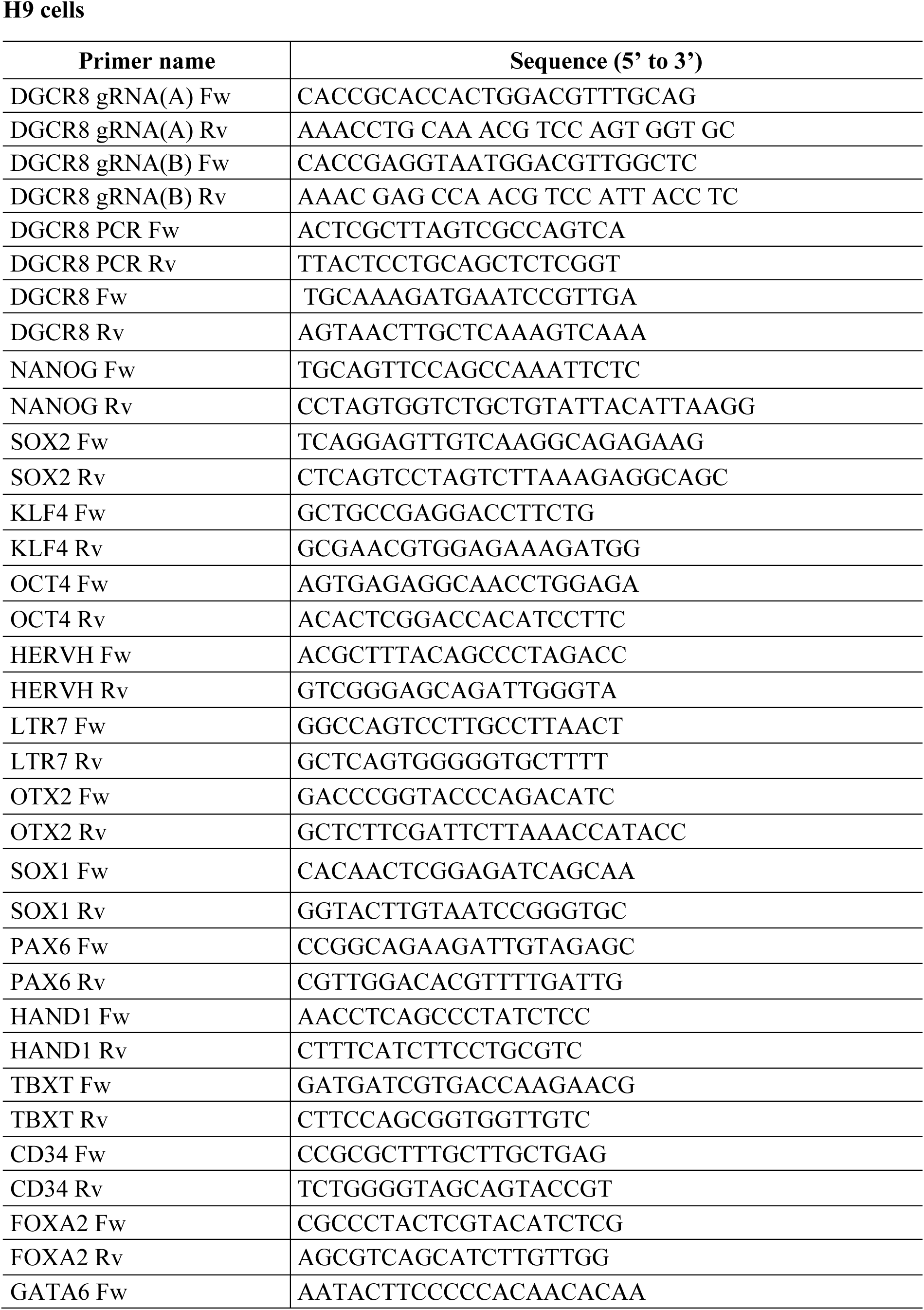
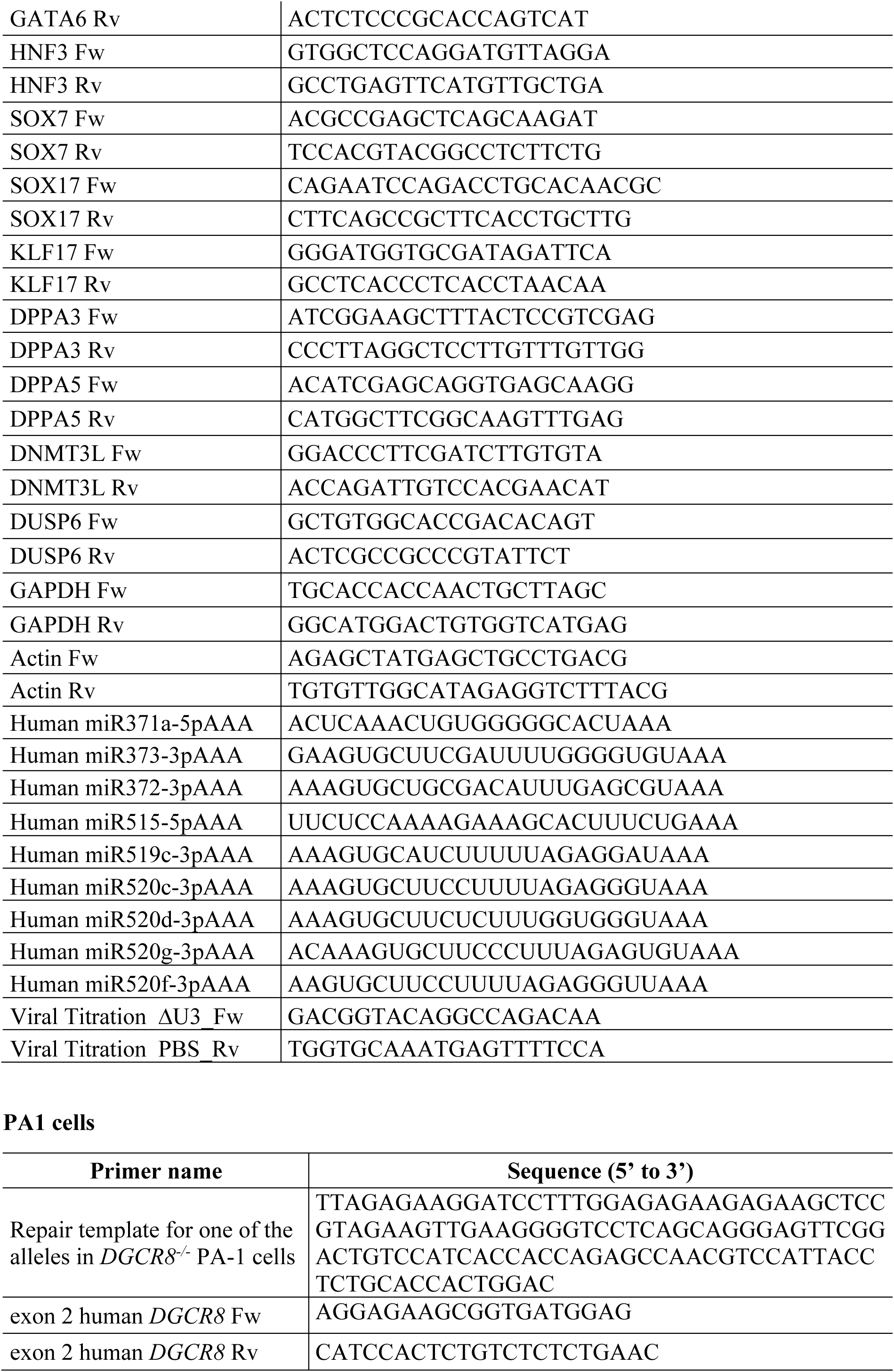
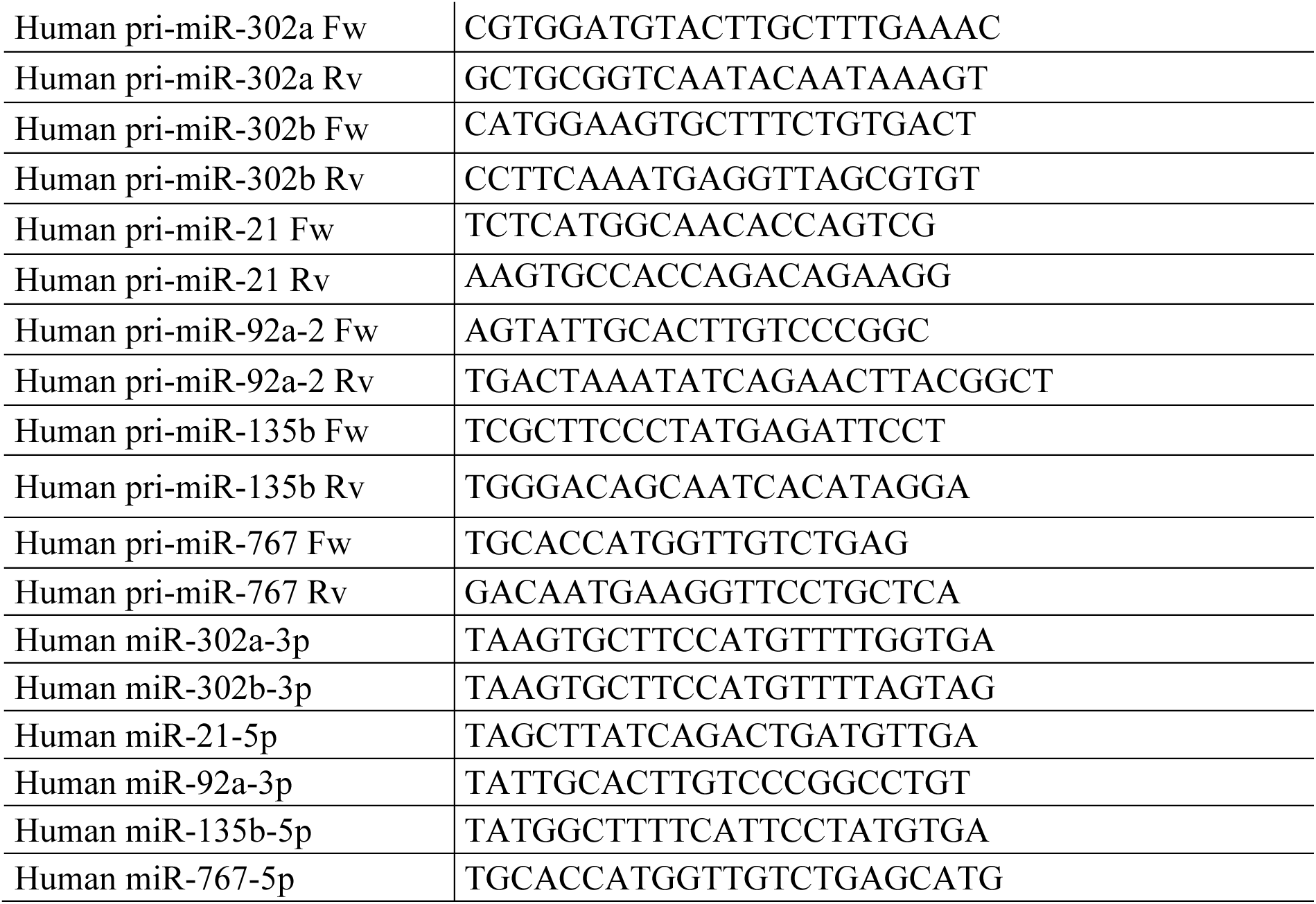
Oligonucleotides used in this study.

### Lentiviral transduction of hESCs

The lentiviral particles overexpressing human DGCR8, pLV-EF1α:hDGCR8 were purchased from Vector Builder. As a lentiviral empty control vector, the promoter EF1α and human *DGCR8* sequences were removed using the restriction enzymes FseI and BsBtI (NEB). After ligation, the right sequence was confirmed by Sanger sequencing. Lentiviral control particles were generated as in (Tristan-Manzano *et al*, 2023). Viral titres (transduction units [TU]/mL) were calculated using qPCR. To this end, 1 x 10^5^ HEK293T cells were transduced with different volumes of the viral supernatant (1, 5 and 10 μl). 72 h post-transduction, genomic DNA was isolated (QiAamp DNA miniKit, Qiagen) and the lentiviral copy number integrated per cell was calculated using a standard curve method. Primers used are listed in **Table 1**. For transduction, hESCs were dissociated and mixed with lentiviruses at a multiplicity of infection (MOI) of 5 and seeded on 24-well plates coated with Matrigel in mTeSR1 medium supplemented with iROCK. After 24 hours, medium was replaced. Three days later, 2 μg/ml puromycin selection was initiated, replacing media every 2 days. After 5 days, transduced hESCs were grown under normal conditions.

### Immunofluorescence

hESCs were seeded on Matrigel-coated coverslips and fixed with 4% paraformaldehyde (PFA) during 5 min followed by permeabilization with 0.1% Triton X-100 in PBS for 5 min at RT. Blocking was performed with 10% donkey serum (Merck Life Science) in PBS containing 0.5% Triton X-100, for 1h at RT. Cells were incubated overnight at 4°C with primary antibodies diluted in PBS containing 1% donkey serum and 0.5% Triton X-100, followed by three washes with 1% donkey serum in PBS. Incubation with secondary antibodies was performed for 30 min at RT, followed by three additional washes with 1% donkey serum in PBS. Cells were counterstained with DAPI (ProLong Gold antifade, Invitrogen) and Zeiss LSM 710 Confocal Microscopy with a Plan-Apochromat 63×/1.40 Oil DIC M27 was used for imaging. Image J was used for quantification. Primary antibodies against Nanog (1:1000, 500-P236, Prepotech), Tra-1-60 (1:100, 09-0068, Stemgent), Brachyury (1:1600, #81694, CST), Sox17 (1:200, AF1924, R&D Systems) and Pax6 (1:200, #60433, CST) were used. Secondary antibodies included an anti-mouse IgG Alexa flour 488 (1:1000, A21202, Invitrogen), anti-rabbit IgG Alexa Fluor 555 (1:1000, A31572, Invitrogen) and anti-goat IgG Alexa Fluor 555 (1:1000, A21432, Invitrogen).

### Western Blot

Cells were lysed in RIPA buffer supplemented with 1x EDTA-free Protease Inhibitor cocktail (Roche), 1mM PMSF and 35 nM β-mercaptoethanol (Sigma). Protein was quantified using the Micro BCA Protein Assay Kit (Thermo Fisher Scientific). Lysates were subjected to SDS-PAGE electrophoresis and transferred to PVDF membranes using Trans-Blot Turbo Transfer System (Bio-Rad). Membranes were blocked using 5% non-fat dried milk in TBS-T (0.1% Tween-20 in 1x TBS) or PBS-T (0.1% Tween-20 in 1x PBS) and incubated with primary and secondary antibodies. Images were acquired using an ImageQuant LAS4000 or Chemidoc Imaging System (Bio-Rad), fluorescent signal was quantified using ImageJ software. Antibodies used include anti-DGCR8 (1:1000, ab90579, Abcam), anti-DROSHA (1:1000, NBP1-03349, NovusBio). anti-SOX2 (1:2000, AB5603, Merck), anti-NANOG (1:500, 500-P236, Peprotech), anti-OCT4 (1:300, sc-8628, Santa Cruz Biotechnology) and anti-KLF4 (1:1000, 4038, Cell Signaling Technology). α-tubulin (1:1000, sc-23948, Santa Cruz Biotechnology) and β-actin (1:15000, A1978, Sigma) antibodies were used as loading controls. As secondary antibodies, anti-rabbit HRP (1:1000, 7074S, Cell Signaling Technology), anti-mouse HRP (1:1000, 7076S, Cell Signaling Technology) and anti-goat HRP (1:10000, 305-035-003, Jackson ImmunoResearch).

### Naïve-like hESCs induction

To revert the primed-state of HET and WT H9 hESCs to a naïve-like state, RSeT™ Feeder-Free Medium (STEMCELL Technologies) was used. Briefly, hESCs were cultured in a T25 flask to 90% confluency. Next, cells were dissociated and detached into aggregates of approximately 100-200 μm in diameter. A dilution (1:20) of the hESCs aggregates was seeded in mTeSR1 medium supplemented with iROCK on Matrigel-coated 6-well plates. One day after seeding, medium was replaced with RSeT™ Feeder-Free Medium and hESCs were grown under hypoxic conditions (37°C, 5% CO_2_ and 5% O_2_). A full-medium change was done every 2 days. After 5 days, hESCs were reverted into naïve-like hESCs.

### Clonal expansion and cell proliferation assays

For clonal expansion assays, 13 x 10^3^ WT, HET(1) and HET(2) H9 hESCs were seeded in 12-well plates coated with Matrigel in mTeSR1 medium. For naïve-like hESCs, 8 x 10^3^ WT, HET (1) and HET (2) were plated in 6-well plates coated with Matrigel in RSeT Feeder-Free Medium. After ten days for hESCs and five days for naïve-like hESCs, cells were fixed (37% formaldehyde, 50% glutaraldehyde in 10× PBS) for 30 min and stained with 0.5 % crystal violet for 40 min at room temperature. Stained areas and number of colonies were quantified using ImageJ software. The area of naïve-like hESCs colonies was measured using cellSens Entry software. For cell proliferation assays, hESCs were seeded at a density of 36,500 cells per well in 12-well plates coated with Matrigel and maintained for 10 days. Cells were counted at days 3, 5, 7 and 10 using a Neubauer chamber. For PA-1 cells, growth rates of WT, HETs and KO cells were compared by seeding 1×10^5^ cells in 6-well plates. Cells were harvested and counted using a hemocytometer before reseeding at day 2, 4 and 7.

### Alkaline Phosphatase staining analysis

H9 hESCs were seeded at a density of 13,000 cells per well in 12-well plate coated with Matrigel for 6 days. Cells were fixed in 4% PFA for 2 min and stained with Alkaline Phosphatase Detection Kit (Sigma-Aldrich). Colonies were manually counted distinguishing between differentiated, mixed and undifferentiated colonies depending on the staining grade and morphology. For single-cells assay, cells were seeded at a density of 0.5 cells/well in a 96-well plate coated with Matrigel and supplemented with cloneR (STEMCELL Technologies) during the first 96 hours after plating. After 10 days, colonies were stained with alkaline phosphatase detection kit. Next, colonies were stained with crystal violet to visualise negative alkaline phosphatase cells.

### Cell-cycle and apoptosis analyses

hESCs were detached using TrypLE and fixed in 70% cold ethanol, washed 3 times with ice cold PBS and centrifuged at 450g for 5 min at 4°C. Cell pellets were resuspended in propidium iodide staining buffer (1x PBS, 0.05% NP-40, 3mM EDTA pH 8, 1 mg/ml RNaseA, 0.05 mg/ml propidium iodide) for 10 min at RT followed by 20 min on ice. Cells were analysed by flow cytometry using FlowJo software. For apoptosis, cells were labelled with the PE Annexin V Apoptosis kit (BD Biosciences) to distinguish between early and late apoptosis by flow-cytometry. Results were represented using BD FACSDiva™ Software.

### H9 hESC differentiation

For EB differentiation, WT and HET H9 hESCs were cultured to 60% confluency. Next, mTeSR1 medium containing Matrigel (1:6 ratio) was added to increase the thickness of the colonies. At 80% confluency, cells were gently detached and cultured in suspension in ultra-low-attachment plates, allowing for EB formation during 21 days in medium, consisting of DMEM Knockout (Gibco) supplemented with 20% FBS (Hyclone), 0.1 mM Non-Essential Amino Acids (Gibco), 1 mM L-glutamine, and 0.1 mM β-mercaptoethanol. Media was replaced every 2 days. For guided differentiations to ectoderm, mesoderm and endoderm lineages STEMdiff Trilineage differentiation kit was used (STEMCELL Technologies). Briefly, H9 hESCs were seeded in mTeSR1 medium supplemented with iROCK on Matrigel-coated coverslips in 24-well plates. 400,000 cells per well were used for ectoderm and endoderm differentiation, and 100,000 for mesoderm. One day after seeding, medium was replaced with lineage-specific medium for 5 days (mesoderm and endoderm), or 7 days (ectoderm). Medium was replaced every day. Cells were next fixed and processed for immunofluorescence analyses as described above.

### RT-qPCR

For RT-qPCR, total RNA was extracted from cells using Trizol, followed by RQ1 DNAse treatment and phenol/chloroform purification. Next, 1µg of total RNA was further treated with DNase I (Invitrogen), and cDNA was synthesised using High-Capacity cDNA Reverse Transcription Kit (Applied Biosystems) and used for qPCR (GoTaq qPCR Mix, Promega). Alternatively, cDNA was synthesised using Transcriptor Universal cDNA Master (Roche) and qPCR was carried out with LightCycler 480 SYBR Green I Master mix (Roche). *GAPDH* or *ACTB* were used as normalisers. Gene expression levels were quantified using the second derivative method. Primers used are listed in **Table 1**.

### miRNA mimics transfection

H9 WT and HETs (7×10^5^) hESCs were seeded in 6-well plates. After 24 hours, cells were transfected with 60 nM of control mimic (4464058, ThermoFisher) or 30 nM of each miR-372-3p and 373-3p mimics (MC10165 and MC11024) or 15 nM of each miR-520g-3p, miR-520d-3p, miR-519c-3p and miR-515-5p mimics (MC10365, MC12807, MH10575 and MC10387,

ThermoFisher) using Lipofectamine 2000 (Life Technologies). 48 hours post-transfection, total RNA was extracted or, alternatively, cells were fixed and stained for clonal expansion assays.

### Chromatin RNA-sequencing and Microprocessor Processing Index

PA-1 cells fractionated similar to (Conrad *et al*., 2014; Witteveldt *et al*., 2018). In brief, 8×10^6^ cells were lysed in mild buffer (10 mM Tris pH 7.4; 150 mM NaCl;0.075% NP-40) and the nuclei and cytoplasm were separated by sucrose gradient. The nucleic fraction was subsequently separated into nucleoplasmic and chromatin-associated fractions as described before, and the chromatin-associated fraction was sonicated on a Bioruptor (5 times 20 sec on/off intervals) prior to DNase treatment using RQ1 DNase (Promega) and RNA extraction using Trizol. Four biological replicate samples for each of the three PA-1 cell lines (WT, HET, KO) were prepared and sent to BGI Genomics for library preparation and high-throughput sequencing after rRNA depletion (2× 100 nt). Reads from chromatin-associated RNA samples were aligned to the human genome (GRCh38.p13) using HISAT2 (v2.1.0) with the options –-no-discordant –-no-mixed –-no-unal (Kim *et al*, 2019). Human pre-miRNA locations were determined by aligning human precursor sequences, obtained using the mature miRNA and hairpin sequences from miRbase (v22.1) against the same genome using Bowtie2 with options –-very-sensitive –-no-unal (Langmead & Salzberg, 2012). The genomic locations of the pre-miRNAs plus 100 nts on each side were determined using bedtools getfasta –s. The read depth for each nucleotide in the alignment of chromatin-associated reads on the appropriate strand for each pre-miRNA was extracted using R.

A 10 nt gap between each pre-miRNA and its flanking regions was created by excluding the outer 5 nts from the pre-miRNA and flanking regions, creating leeway for potential alternative Drosha cleaving. For each region, the average read depth of all included nucleotides was used for Microprocessor Processing Index (MPI) calculation. MPI was defined as the negative log2-transformed ratio between the mean read depth (RD) of the pre-miRNA region (hairpin) to the mean read depth of the regions flanking the pre-miRNA. In this manner, a high MPI and > 0 indicates efficient processing, and a MPI close to 0 or negative, indicates inefficient or absence of processing.

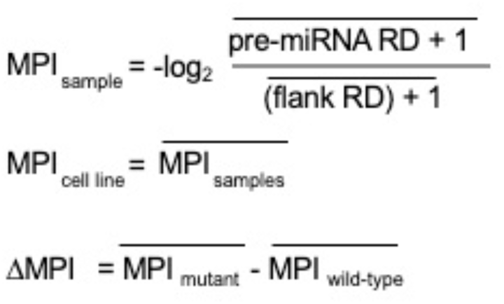

To filter out Microprocessor-independent pri-miRNAs, reduce noise and exclude artifacts, pri-miRNAs were only included if they: (1) were no mirtrons or tailed mirtrons; (2) produce miRNAs which were detected and included in statistical analysis for small RNA sequencing; (3) on average, the flank depth in WT samples is ≥ 2; (4) the ratio between the two flanks did not differ more than 4-fold. Differences in processing for individual pri-miRNA between cell lines (ΔMPI) were computed by subtracting MPI values in *DGCR8*(WT) vs (HET) or (KO) cells (ΔMPI= MPI(HET/KO) – MPI(WT)). A negative ΔMPI indicates that the pri-miRNA is less processed in HET/KO compared to WT cells.

### MiRNA quantification and small RNA high-throughput sequencing

MiRNA quantification in H9 hESCs was performed as described (Tristán-Ramos *et al*, 2020). For PA-1 cells, 200 ng of total RNA was used for the retrotranscription (RT) reaction using the miRCURY LNA miRNA PCR system (Qiagen). After dilution of the resulting cDNA, qPCR was performed using the miRCURY LNA SYBR Green PCR Kit (Qiagen) with corresponding miRNA primers on a LightCycler 480 Instrument (Roche). Both H9 and PA-1 miRNA quantification data were normalised to the DGCR8-independent miRNA, *hsa-miR-320a*. For miRNA-specific primers see **Table 1**.

For small RNA high-throughput sequencing, RNA from three different biological replicates for WT, HET(1) and HET(2) hESCs was extracted using mirVana microRNA isolation kit (ThermoFisher). Small RNA libraries were generated using NEXTFlex Small RNA Library Prep Kit v3 (Cat #NOVA-5132–06) and sequenced on the NextSeq 500 system (Illumina, CA, USA) by the Genomic Unit at Genyo. For PA-1 cells, RNA from four different biological replicates from WT and HET cells was extracted using the miRNeasy Mini Kit (Qiagen) followed by on-column DNAse digestion. Small RNA sequencing libraries using unique molecular identifiers (UMIs) were sequenced using DNB-seq (Li *et al*, 2019) by BGI. For each sample, identical small RNA-seq reads were collapsed and only reads that were present more than once were included in further analysis. MiRNAs were identified and counted using miRDeep2, using the quantification function with human hairpin and mature miRNA sequences files from miRbase (v22.1) and allowing no mismatches (Friedlander *et al*, 2012; Kozomara *et al*, 2019; Kozomara & Griffiths-Jones, 2014). DESeq2 was used for statistical analysis, comparing WT vs HET in H9 and PA-1 separately, and using apeglm as log fold-change shrinkage model (Love *et al*, 2014; Zhu *et al*, 2019).

### *In silico* miRNA target prediction and pathway analysis

Differentially expressed miRNAs (p-adj ≤ 0.05) that are common to both H9 hESCs HET(1) and HET(2) (85 miRNAs) and differentially expressed miRNAs in PA-1 HET (155miRNAs) underwent functional enrichment analysis with DIANA-mirPATH v3 software (Vlachos *et al*, 2015) (for PA-1 HET, only the top 100 miRNAs with the highest absolute log2FC were used). MicroT-CDS prediction algorithm was used to identify putative mRNA targets of miRNAs and associated significantly enriched (FDR ≤ 0.05) KEGG pathways (Kanehisa *et al*, 2014) identified. Dot plots were generated using ggplot2 (v3.3.5) and DOSE v3.14.0 (Yu *et al*, 2015) R packages.

### Total RNA high-throughput sequencing and gene ontology analyses

Total RNA from three biological replicates of H9 (WT, HET(1) and HET(2)) and four biological replicates of PA-1 (WT and HET) cells was extracted using miRNeasy Mini Kit (Qiagen) followed by on-column DNAse digestion. Purified RNA was rRNA-depleted prior to sequencing (DNB-seq) by BGI. For PA-1 RNA-seq analyses, paired-end reads were aligned to the human genome (GRCh38.p13) using HISAT2 with options –-no-discordant –-no-mixed –-no-unal (Kim *et al*., 2019). Transcript counts for each sample were created with featureCounts (Rsubread v2.2.6), using reverse counts and excluding reads that aligned to the genome multiple times (Liao *et al*, 2014). For H9 RNA-seq analyses, quality of individual H9 WT and HET sequences were evaluated using FastQC v0.11.5 software (Andrews, 2010). Paired-end reads were aligned to GRCh38.p13 human genome assembly with STAR v2.7.6a (Dobin *et al*, 2013) and quantified with featureCounts v2.0.1 (Liao *et al*., 2014) using NCBI Annotation Release 109. For both H9 and PA-1, differential expression analysis and count normalization was performed with the R package DESeq2 v1.28 (Love *et al*., 2014). After differential analysis, DESeq2’s apeglm lfcShrink was applied to shrink log_2_ fold changes (Love *et al*., 2014; Zhu *et al*., 2019).

Functional enrichment analysis was carried out for differentially expressed genes (p-adj ≤ 0.05) using the enrichGO function in R package clusterProfiler v3.16.1 (Yu *et al*, 2012), and using the Gene Ontology Biological Processes (GO-BP) gene sets.

### Analysis of primate-specific C19MC cluster

A list of primate-specific miRNAs was obtained from MirGeneDB 2.1(Fromm *et al*., 2022). R package miRBaseConverter v1.14.0 (Haunsberger *et al*, 2017) was used to adapt this list to mirBase v22.1 used for differential expression analysis. All miRNAs with padj ≤ 0.05 were considered as dysregulated. Common primate-specific dysregulated miRNAs in H9 HET (34 miRNAs) and PA-1 HET cells (47 miRNAs) were analysed using DIANA-mirPATH v3 software. Statistical significance for the enrichment of primate-specific dysregulated miRNAs was calculated using Fisher’s exact test (phyper function of R package stats, v4.2.1).

### Transposable element expression analyses

The SQuIRE v0.9.9.92 pipeline (Yang *et al*., 2019) was used to measure transposable elements expression changes using default parameters. SQuIRE downloads TE annotations from RepeatMasker and uses an expectation-maximization algorithm to assign multimapping reads. Next, it performs TE differential expression using DESeq2, either by grouped TE subfamilies or by analysing individual loci. Expression of individual TEs was represented using ggplot2 (v3.3.5) and gghalves (v0.1.1) (Tiedemann, 2020) R packages. hESCs-specific chimeric transcripts and lncRNAs derived from HERVH were extracted from (Wang *et al*., 2014) and all gene symbols were updated using the R package HGNChelper v0.8.1 (Oh *et al*, 2020). This gene list was compared with differentially expressed genes (p-adj ≤ 0.05) from hESCs HET(1) and HET(2). Heatmap was generated using the R package ComplexHeatmap v2.4.3(Gu *et al*, 2016) applied to Z-score for each gene and volcano graphs using ggplot2 (v3.3.5). Normalized bigwig files were generated using Draw tool from SQuIRE and visualized using IGV.

### Assay for Transposase Accessible Chromatin with sequencing (ATAC-seq)

ATAC-seq libraries were prepared as previously described (Buenrostro *et al*, 2015). Around 50,000 cells were used for each biological replicate (n=3). Cells were lysed in 50 µL cold lysis buffer (10 mM Tris-HCl pH 7.4, 10 mM NaCl, 3 mM MgCl2 and 0.1% IGEPAL CA-360) and pelleted at 500g for 10 mins at 4°C. Pellets were resuspended in 50 µL transposition reaction mix as follows: 2X Tagment DNA buffer (Illumina 15027866), 20X Tagment DNA enzyme (Illumina 15027865) and incubated at 37°C for 30 mins. Transposed samples were purified using the MinElute PCR purification kit (Qiagen 28204), and eluted in 10 µL. Transposed DNA samples were amplified by PCR by setting up a 50 µL reaction as follows: 10 µL transposed DNA, 9.7 µL ddH2O, 2.5 µL 25 µM customised Nextera PCR primer FW, 2.5 µL 25 µM customised Nextera PCR primer RV, 0.3 µL 100X SYBR Green I (Invitrogen S-7563) and 25 µL 2X NEBNext high-fidelity PCR master mix (NEB, M0541). To reduce GC/size bias and over amplification of libraries, the reaction was monitored by qPCR to stop amplification prior to saturation. A qPCR 15 µL side-reaction was set up as follows: 5 µL 5 cycles PCR amplified DNA, 4.44 µL ddH2O, 0.25 µL 25 µM customised Nextera PCR primer FW, 0.25 µL 25 µM customised Nextera PCR primer RV, 0.06 µL 100X SYBR Green I and 5 µL 2X NEBNext high-fidelity PCR master mix. The additional number of cycles required for each sample was determined by plotting a linear run vs. the cycle number. The number of cycles that correspond to ¼ maximum fluorescent intensity was calculated for each sample. The remaining 45 µL 5 cycles PCR amplified DNA was run as before (omitting 72°C initial step and modifying the number of cycles to the calculated amount). Amplified DNA samples were subjected to double size selection to remove DNA fragments <150 base pairs (bp) and >1000 bp that would hinder sequencing reactions. Samples were diluted up to 90 µL with ddH2O and purified using the SPRIselect beads (Beckman Coulter B23317). For removal of large DNA fragments, a ratio of 0.55 DNA/beads slurry was used, and for removal of small DNA fragments, a ratio of 1.8 was used. Purified DNA samples were quantified using the Qubit dsDNA high sensitivity assay (ThermoFisher Q32854) with the Qubit 4 fluorometer. The quality of DNA samples was analysed using the high sensitivity DNA kit (Agilent 5067-4626) in Bioanalyzer. Samples with the nucleosomal fragment distribution profile expected for ATAC-seq libraries (mono-, di-and trisomal fragments) were sent for sequencing.

Samples were sequenced on an Illumina HiSeq 4000 platform to obtain 50 bp paired-end reads at the Wellcome Trust Clinical Research Facility (University of Edinburgh). Reads were trimmed using cutadapt v3.5 paired-end trimming, and aligned to the hg38 human genome using bowtie2 v2.4.4 (Langmead & Salzberg, 2012) paired-end alignment with options –-very-sensitive –-no-mixed –-no-discordant –X 2000. Unmapped reads and those mapping to the mitochondrial genome were removed and duplicate reads were filtered out using Picard 2.27.5 MarkDuplicates (http://broadinstitute.github.io/picard/). Reads were shifted by +4 bp for those mapping to the positive strand and −5 bp for those mapping to the negative strand using alignmentSieve tool from deepTools package. Broad peaks were called using MACS2 2.2.7.1 callpeak (Zhang *et al*, 2008) with options –g hs –f BED –-keep-dup all –q 0.01 –-nomodel –shift –75 –-extsize 150 –-broad. Peaks overlapping blacklisted regions were removed using bedtools. A union peak set across all samples was obtained following the iterative overlap peak merging procedure described on (Grandi *et al*, 2022) (code provided in https://github.com/corceslab/ATAC_IterativeOverlapPeakMerging). A count matrix over the union peak set was computed using featureCounts 2.0.1 (Liao *et al*., 2014) and differently expressed peaks were obtained using DESeq2 1.28 R package (Love *et al*., 2014) and selecting dysregulated peaks using p-adj ≤ 0.05 and abs(log2FC) ≥ 1. Peaks were annotated using annotatePeaks.pl from Homer v4.11 package (Heinz *et al*, 2010). Functional analysis of differential ATAC-seq regions was carried out with rGREAT package (Gu & Hubschmann, 2023). Motif analysis was performed with findMotifsGenome.pl tool from Homer v4.11 package using default parameters and random background selection. Metagene plots were generated using computeMatrix and plotProfile tools from deepTools. Genes contained in an interval of ± 10kb from differentially expressed PA1 HET ATAC-seq peaks were obtained using annotatePeakInBatch function in ChIPpeakAnno R package(Zhu *et al*, 2010).

## Data and code availability

All PA-1 sequencing data are deposited in GEO database, accession number GSE197474 (token: *qbulgmakldgrbap*). ATAC-seq datasets are deposited in GEO, accession number GSE205798 (token: *ivaduicidhwnlor*). H9 data are deposited in GEO, accession number GSE209843 (token: *obsnscqwbfshfuz*). Bioinformatic and software packages are described in the STAR Methods sections.

## Acknowledgements

The work in the lab of S.R.H was supported by Grant PID2020-115033RB-I00 funded by MCIN/AEI/10.13039/501100011033, grants PEJ2018-003280-A and RYC-2016-21395 funded both by AEI /10.13039/501100011033 and by “ESF investing in your future”, Career Integration Grant-Marie Curie (FP7-PEOPLE-2011-CIG-303812), FEDER/Junta de Andalucía-Consejería de Transformación Económica, Industria, Conocimiento y Universidades (PY20_00619 y A-CTS-28_UGR20 grants) and donation to “Aula de estudios 22qDS”. The work in S.M laboratory was funded by the Wellcome Trust grants (221737/Z/20/Z and 107665/Z/15/Z), the Royal Society grant (RGS\R1\191368) and the Wellcome Trust iTPA (PIII021). L.K. was funded by an MRC-Precision Medicine fellowship and P.C was funded by a Darwin Trust fellowship. A.G.-G was supported by grant PRE2021-098878 funded by MCIN/AEI/10.13039/501100011033. J.L.G.-P. acknowledges funding from ERC (ERC-Consolidator ERC-STG-2012-309433), the Government of Spain (MINECO-FEDER SAF2017-89745-R and PID2021-128934NB-I00), the Andalusian regional Government (PAIDI P12-CTS-2256 and P18-RT-5067) and a private donation from Ms Francisca Serrano (Trading y Bolsa para Torpes, Granada, Spain). We are grateful to Javier F. Caceres for comments and critical reading of the manuscript. We thank Gregorz Kudla for initial bioinformatic analysis. Paul de Sousa and Rosa Montes for advice on hESC biology and techniques; Marta García-Canadas, Jennifer Parra, Esther Prada, Meriem Benkaddour-Boumzaouad and the Genyo’s microscopy unit for technical support. We thank the Francisco Martin’s group (Genyo, Granada) for help with the lentiviral transduction experiments. We would like to specially thank the Associations of 22q11.2 patients in Andalucia and Levante for their support and trust.

## Author contributions

S.M. and S.R.H. conceived and supervised this study. A.C.B., L.K., S.M. and S.R.H designed and interpreted the experiments. A.C.B. performed most experiments with hESC. L.K. performed most experiments with PA-1 cells and performed data analyses for transcriptomic approaches, under the supervision of A.I. and S.M. G.P. provided additional data analysis for hESC and PA-1 datasets. P.C. and K.G. performed experimental validation with PA-1 cells. L.S. provided technical support and performed experiments. S.P. performed and analysed ATAC-seq experiment in the group of R.H. P.T.R generated DGCR8 knockout PA1 cells and DGCR8 heterozygous hESCs. A.G.G. contributed to HERVH analysis. J.L.G.P contributed with resources to generate DGCR8 heterozygous hESCs. G.B. performed preliminary bioinformatic analysis. S.M. and S.R.H wrote the original draft and all the authors contributed to the final version.

## Figure legends

**EVF 1. Characterization of *DGCR8* heterozygous human pluripotent cell models**.

(**A**) Sanger sequencing of H9 hESC *DGCR8* HET clone 1 (left) and clone 2 (right). HET(1) contains a 35-nt insertion in one allele of *DGCR8* that contains a stop codon. HET(2) contains a frameshift 8-nt deletion in one allele of *DGCR8* (**B**) Sanger sequencing of *DGCR8* KO PA-1 cells, allele 1 contains a 60 nt insertion which includes a stop codon, and allele 2 contains a frameshift 7-nt insertion in exon 2 (**C**) Sanger sequencing of *DGCR8* PA-1 HET clones (both HET1 and HET2 are genetically identical). HET cells were generated by correcting the frameshift 7-nt insertion in exon 2 of *DGCR8* in KO cells (**D**) DGCR8 and DROSHA western blot analyses of WT, two different HET DGCR8 PA-1 and KO clones. α-Tubulin serves as a loading control (**E**) Quantification of DGCR8 and DROSHA protein levels in PA-1 WT, HET and KO cells. Data represent the average of three independent experiments +/− st.dev. α-Tubulin serves as a loading control (**F**) DGCR8, DROSHA and KLF4 western blot analyses of H9 WT and HET DGCR8 hESCs transduced with a control or a lentiviral vector expressing human *DGCR8*. Actin serve as loading control (**G**) RT-qPCR analyses of KLF4 for WT and HET hESCs transduced with control or lentiviral vector expressing human *DGCR8*. Data are normalised to *GAPDH* and relative to WT levels. Data are the average (n=3) +/− st. dev. (***) p-val ≤ 0.001, (****) p-val ≤ 0.0001, by one-way ANOVA followed by Tukey’s multiple comparison test.

**EVF 2. *DGCR8* heterozygous human pluripotent cell models display proliferation and pluripotent defects.**

(**A**) Alkaline phosphatase staining of WT and HET H9 hESCs, with high (left) or low (right) magnification. Quantification of alkaline positive colonies, distinguishing between positive or mixed, regarding morphology and staining intensity (**B**) Quantification of alkaline positive, mixed or negative (differentiated) colonies, obtained after single-cell dilution of WT and HET cells. Data are the average of three independent experiments +/− st. dev., (***) p-val ≤ 0.001 by two-way ANOVA, followed by Dunnett’s multiple comparison test (**C**) Clonal expansion assay for WT and HET cells transduced with a control or a lentiviral vector expressing human *DGCR8*. Colonies are visualised by crystal violet staining (**D**) Cell proliferation essay of WT and HET hESCs transduced with control and DGCR8-expressing lentiviruses (left) and PA-1 WT, HET and KO cells (right) (**E)** Cell cycle analyses by flow cytometry. Data represent the average of 3 biological replicates +/− st. dev. (*) p-val ≤ 0.05, (**) p-val ≤ 0.01 by two-way ANOVA followed by Dunnett’s multiple comparison test (**F**) Representative flow cytometry scatter plots for WT and HET hESCs transduced with a control lentiviral vector or DGCR8-expressing vector, stained with 7-AAD (y-axis) and PE Annexin V (x-axis). Cells positive for PE Annexin V are designated as ‘early apoptosis’, double labelled are considered ‘late apoptosis’.

**EVF 3. Functional analysis of primate-specific dysregulated miRNAs in both human pluripotent cell models**.

(**A**) Fraction of expressed and significantly dysregulated primate-specific miRNAs (pink). Dysregulated miRNAs were significantly enriched in primate-specific miRNAs in all HET cell lines: HET1 H9 hESCs (p-val =6.835985e-05), HET2 H9 hESCs (p-val =2.488492e-04) and HET PA-1 (p-val =3.65392e-07) (**B**) KEGG pathway analyses for the predicted targets (microT-CDS) of all the common significantly dysregulated miRNAs in HET DGCR8 hESCs (**C**) Volcano plot for miRNA expression in HET PA-1 cells vs WT. MiRNAs in pink are primate-specific miRNAs (**D**) KEGG pathway analyses for predicted targets (microT-CDS) of dysregulated miRNAs (TOP100) in HET PA-1 cells (**E**) the same as (**D**), but only using primate-specific miRNA predicted targets. Common pathways in (**E**) and (**D**) are in black, unique in grey.

**EVF 4. Analysis of gene expression defects in both human pluripotent cell models**.

(**A**) Volcano plots for differentially expressed genes in the two HET hESC clones vs WT hESCs. Blue are downregulated genes, and red, upregulated (**B**) the same as in (**A**) for HET PA-1 cells (**C**) GO pathway enrichment for differentially expressed genes in *DGCR8* HET PA-1 cells (only included TOP 30 categories) (**D-E**) Kernel density estimation of log2FC distributions for predicted targets of all dysregulated miRNAs (blue), targets of primate-specific dysregulated miRNAs (pink) and controls or non-target genes (black) for (**D**) PA-1 HET cells and (**E**) HET (1) and HET (2) hESCs. Distribution is significantly different for both targets of all dysregulated miRNAs and primate-specific miRNAs vs. non-targets (p-val < 2.22e-16).

**EVF 5. *In vitro* pri-miRNA processing, chromatin-associated RNA enrichment and ATAC-seq controls**

(**A**) Top, Ponceau staining of cytoplasmic (cyto), nucleoplasmic (nucl) and chromatin (chr) fractions from WT, HET and KO PA-1 cells. Bottom, western blot analyses of the same fractions against TUBULIN, which serves as a cytoplasmic fraction marker, and histone H3, which is a chromatin marker (**B**) RT-qPCR analysis of cytoplasmic, nuclear and chromatin fractions. *GAPDH* pre-mRNA serves as a positive control for chromatin fractions, while *GAPDH* mRNA is enriched in the cytoplasm. All three tested pri-miRNAs were enriched in the chromatin fractions from WT PA-1 cells. An equal proportional volume from the three fractions was used for cDNA preparation and qPCR analyses. Data for each primer pair are represented as a fold-change over the ‘cytoplasmic’ sample (**C**) Read depth coverage across several DGCR8-dependent miRNAs from chromatin-associated RNA high-throughput sequencing in WT, HET and KO PA-1 cells. Grey boxes indicate mature miRNAs, black line represents surrounding genomic regions (**D**) Read distribution around TSS (transcription start sites) of ATAC-seq peaks for WT (blue), and HET (orange) PA-1 cell lines.

