## Supplementary figures and images for "*DGCR8* haploinsufficiency leads to primate-specific RNA dysregulation and pluripotency defects"

### Supplemental Figure 1

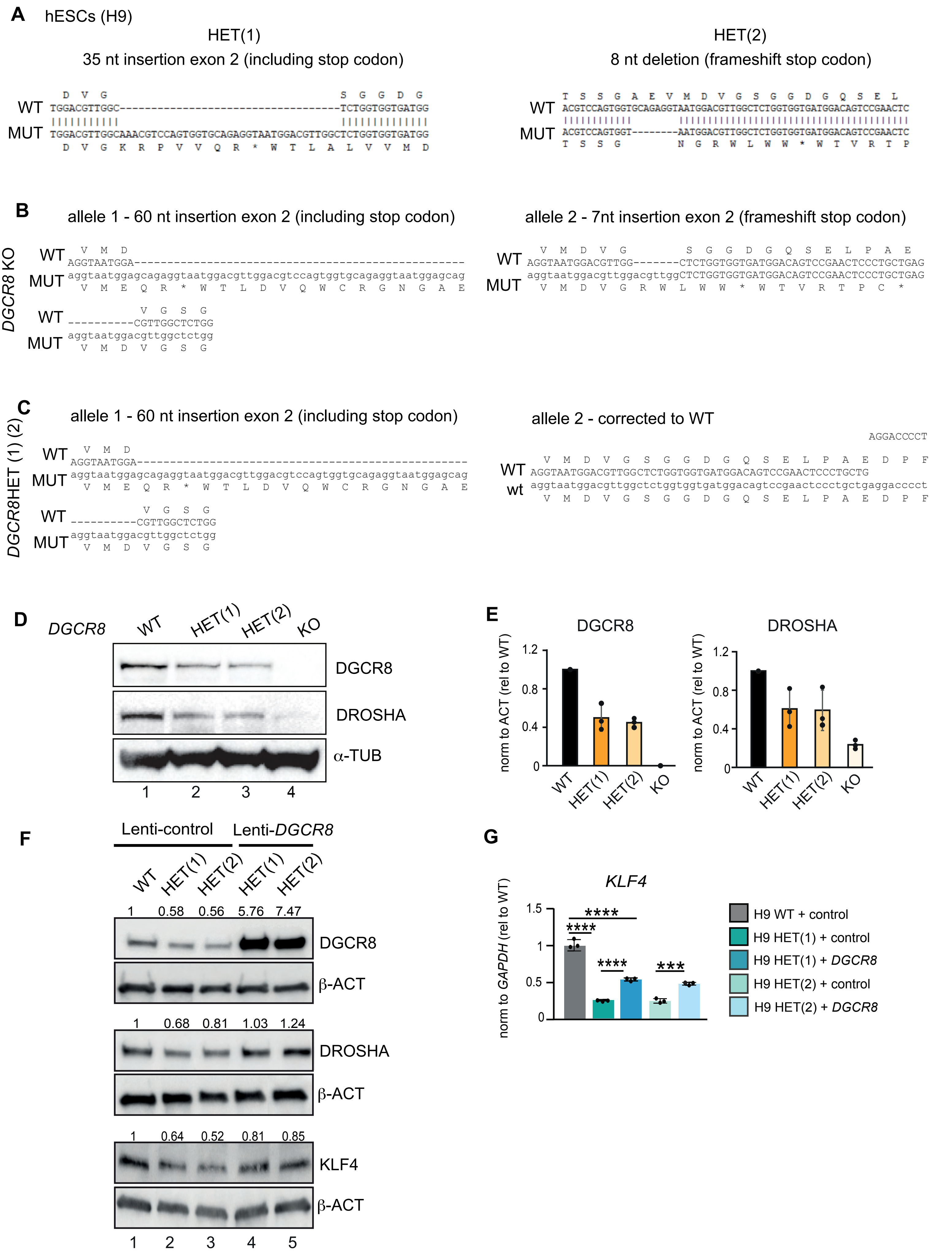

### Supplemental Figure 2

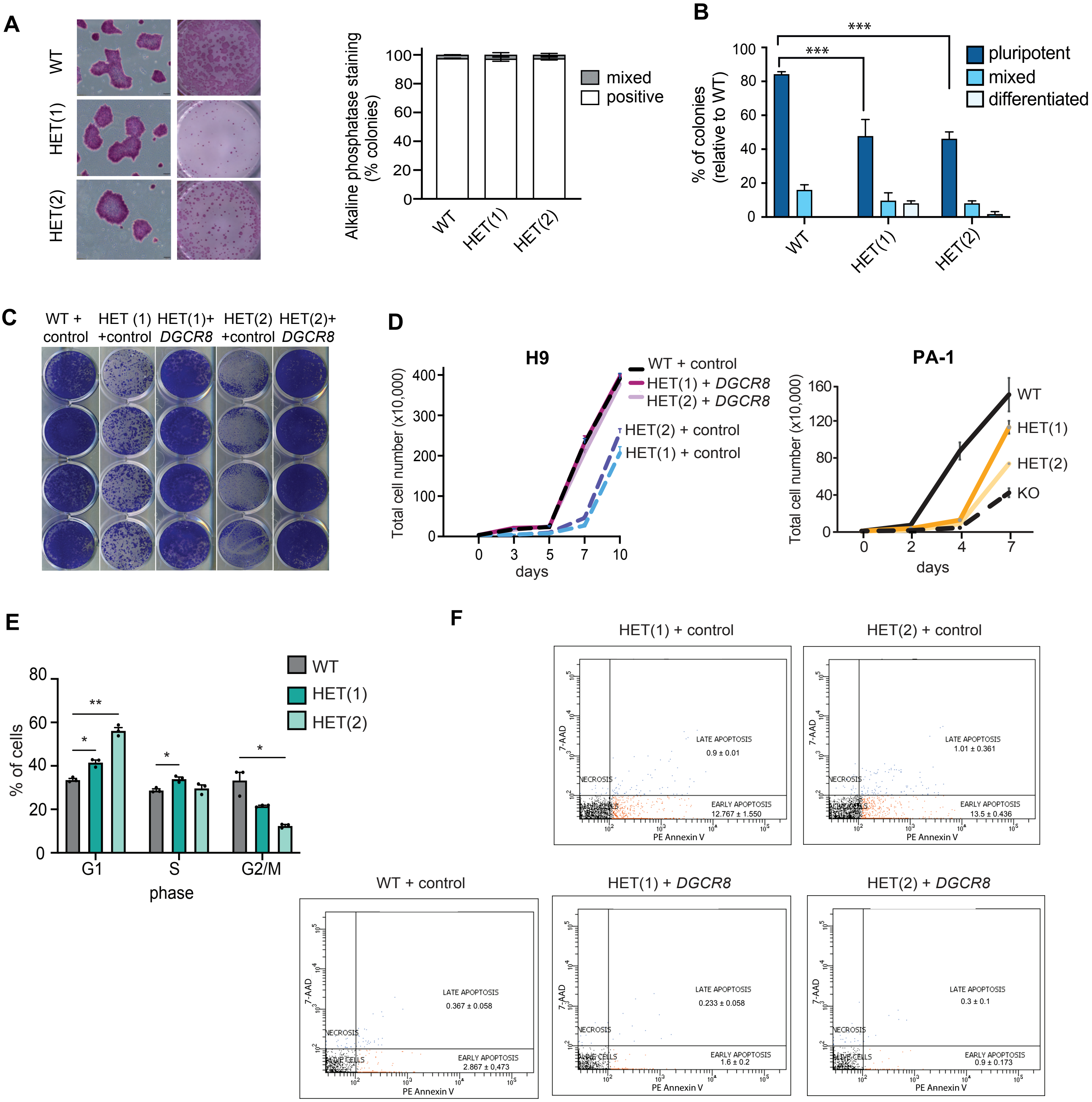

### Supplemental Figure 3

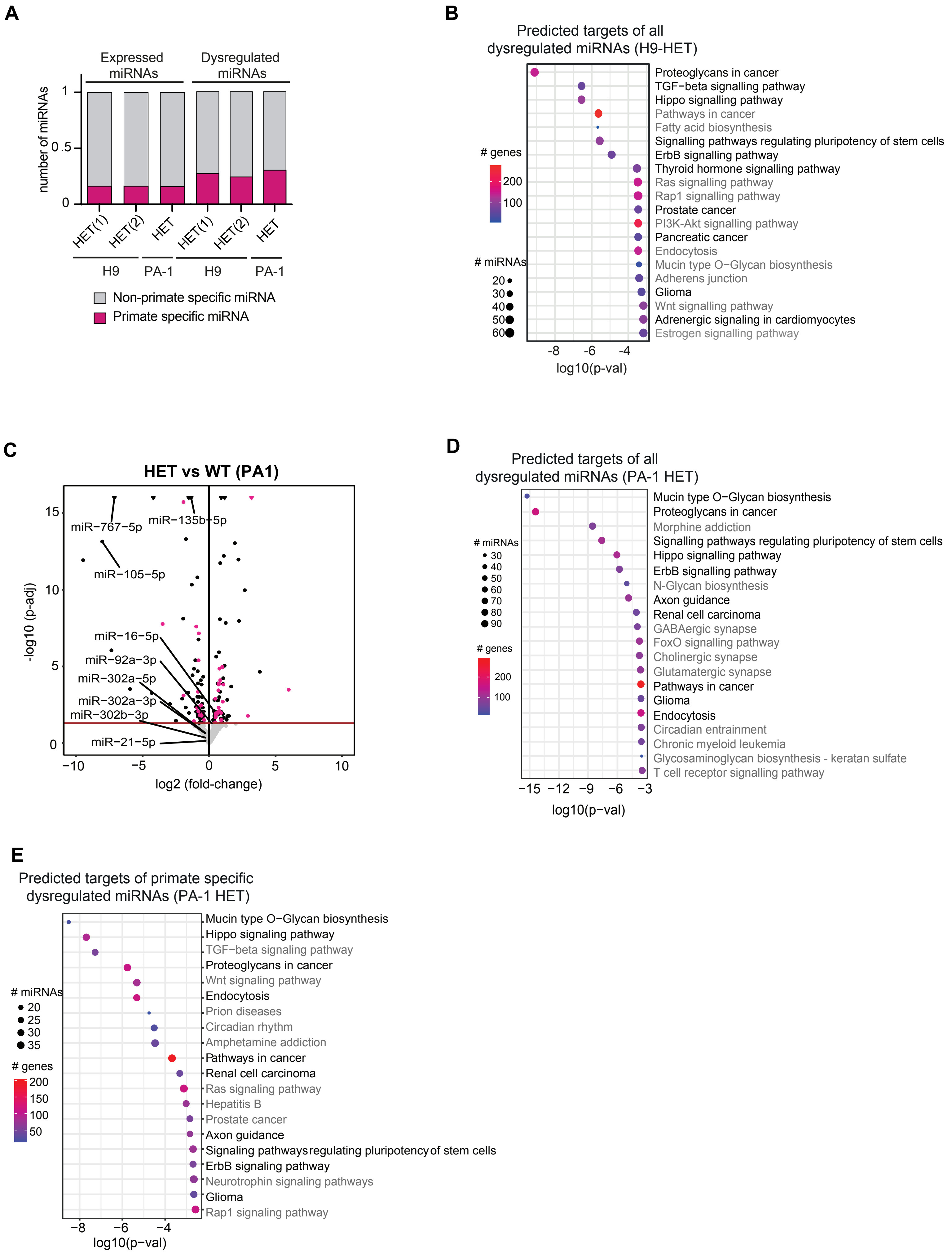

### Supplemental Figure 4

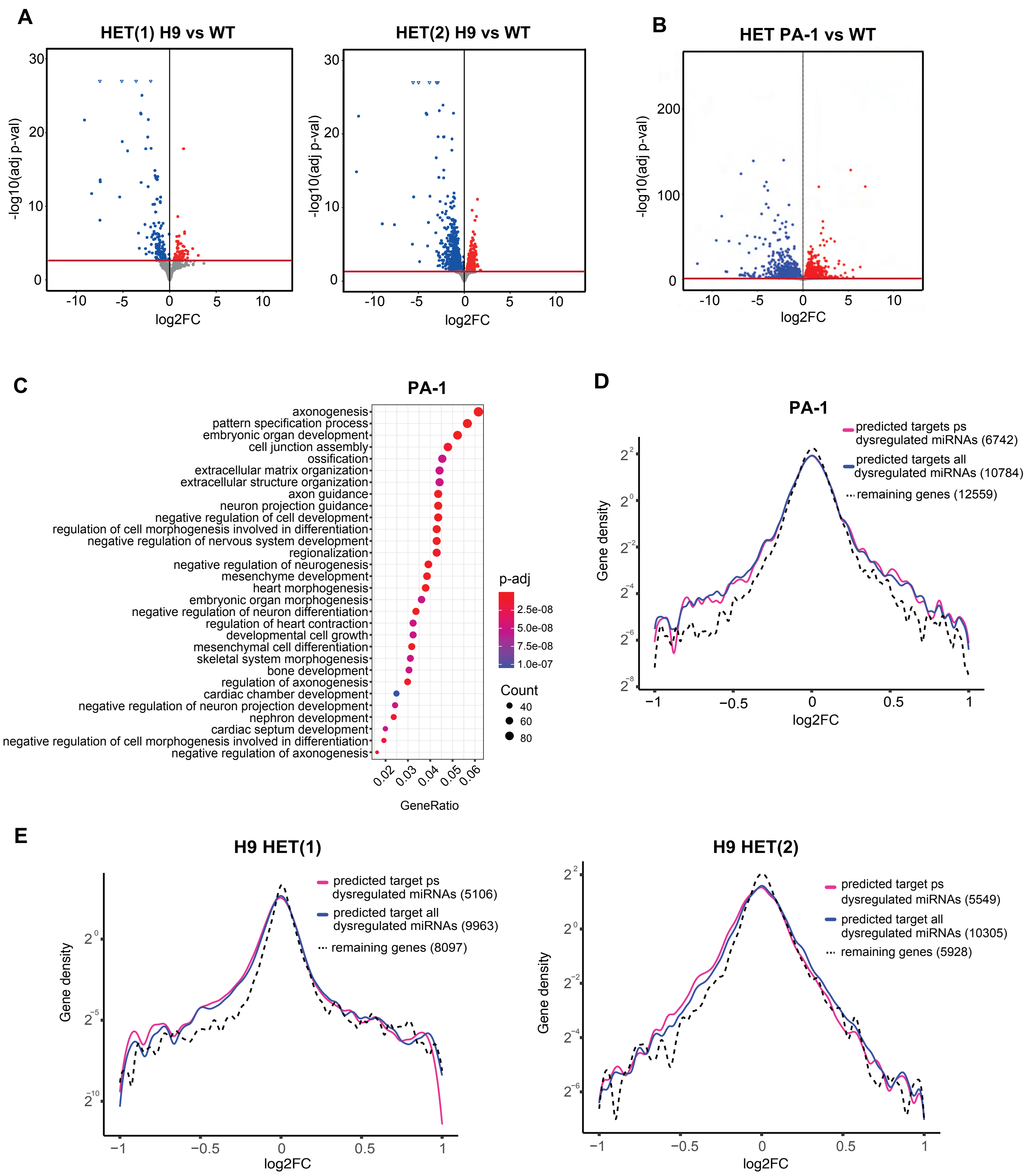

### Supplemental Figure 5

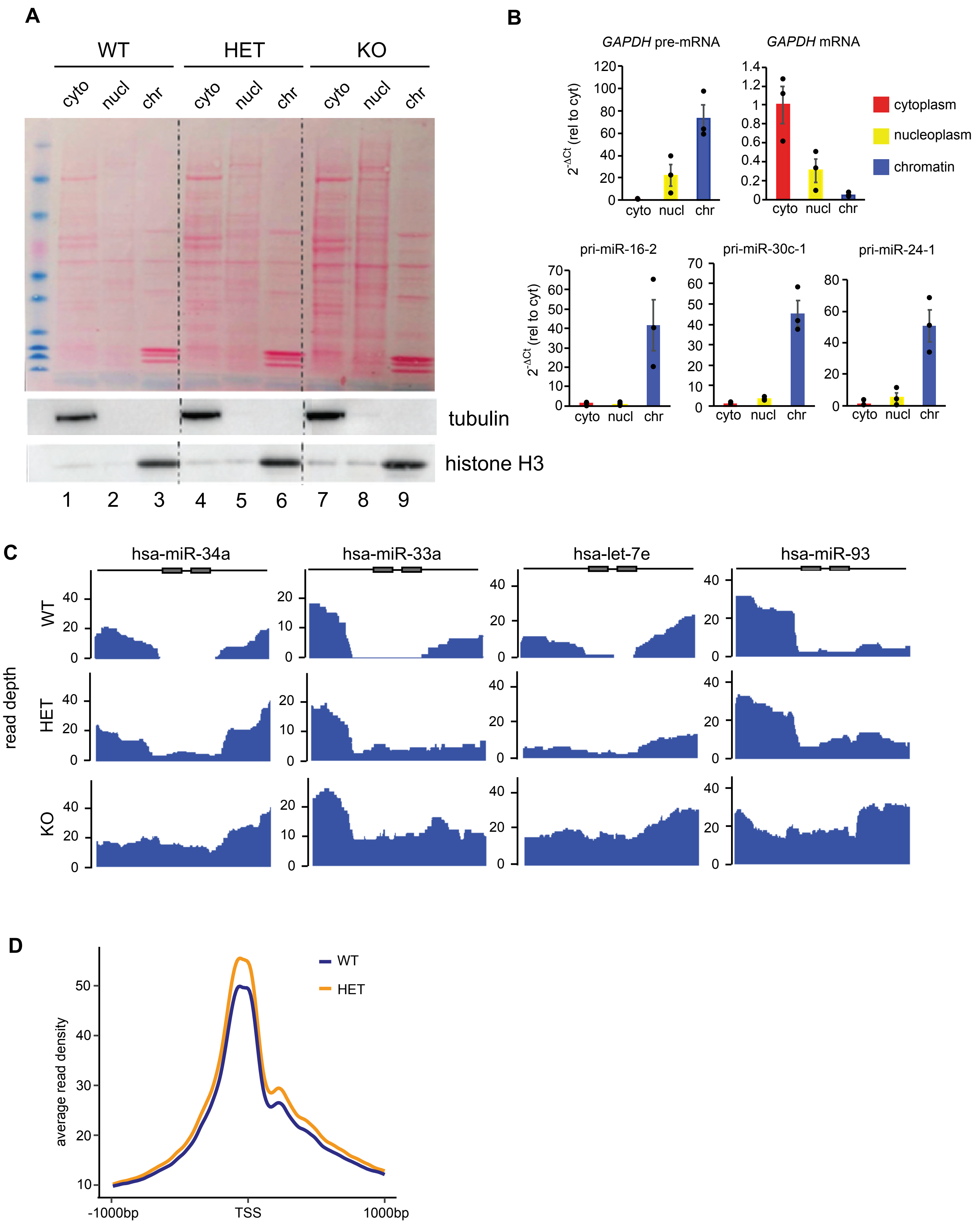
